# Yorkie controls tube length and apical barrier integrity in the developing *Drosophila* airways

**DOI:** 10.1101/532325

**Authors:** Dimitrios K. Papadopoulos, Pavel Tomancak, Vasilios Tsarouhas, Christos Samakovlis, Elisabeth Knust, Kassiani Skouloudaki

## Abstract

Epithelial organ size and shape depend on cell shape changes, cell-matrix communication and apical membrane growth. The *Drosophila* embryonic tracheal network is an excellent model to study these processes. Here, we show that the transcriptional co-activator of the Hippo pathway, Yorkie (YAP in vertebrates), plays distinct roles in the developing *Drosophila* airways. Yorkie exerts a cytoplasmic function by binding *Drosophila* Twinstar, the orthologue of the vertebrate actin-severing protein Cofilin, to regulate F-actin levels and apical cell membrane size, which are required for proper tracheal tube elongation. Second, Yorkie controls water-tightness of tracheal tubes by transcriptional regulation of the enzyme *δ-aminolevulinate synthase* (*Alas*). We conclude that Yorkie has a dual role in tracheal development to ensure proper tracheal growth and functionality.

**Short Summary:** This work identified an alternative role of the transcriptional co-activator Yorkie (Yki) in controlling water impermeability and tube size of the developing *Drosophila* airways. Tracheal impermeability is triggered by Yki-mediated transcriptional regulation of *δ-aminolevulinate synthase*, *Alas*, whereas tube elongation is controlled by binding of Yki to the actin severing factor Twinstar.

## Introduction

Regulation of epithelial tube size and integrity depends on various control mechanisms, including cell surface receptors, cytoskeletal and extracellular matrix components, cell polarity and vesicular transport. These distinct cellular mechanisms control the development and maintenance of tube diameter and length, which are key requirements for proper tube function (Beitel and Krasnow, 2000; Iruela-Arispe and Beitel, 2013). Aberrant regulation of any of these components causes human diseases, such as polycystic kidney disease (PKD) (Steinman, 2012), fibrocystic breast disease (Rinaldi et al., 2010), pancreatic cystic neoplasma (Garud and Willingham, 2012) or thyroid nodules (Popoveniuc and Jonklaas, 2012). To understand the biology of epithelial tube formation and functionality, several models have been used, including the well-characterized *Drosophila* respiratory system, the tracheae. The tracheae form a branched tubular network during the first half of embryogenesis. At mid-embryogenesis, the tubes expand and a transient cable is deposited in the tubular lumen, consisting of a chitinous apical extracellular matrix (aECM), which confers rigidity to the tubes and is responsible for diametric expansion and axial elongation (Forster et al., 2010; Luschnig et al., 2006; Tonning et al., 2005; Wang et al., 2006). Later, endocytosis removes the luminal proteins and the tubes fill with air (Hayashi and Dong, 2017; Tsarouhas et al., 2007). During late embryonic development and after diametrical expansion, tracheal tubes elongate by means of cell shape changes and cell-cell junction rearrangements, but without increasing their cell number (Samakovlis et al., 1996). So far, it has been well established that longitudinal growth depends on various cellular components and molecules, including: i) septate junctions (Llimargas et al., 2004; Wang et al., 2006; Wu et al., 2004), ii) the subapical protein Crumbs and the apical cytoskeleton (Dong et al., 2014; Laprise et al., 2006; Laprise et al., 2010; Tonning et al., 2005), and iii) chitin deacetylases and Src kinase levels (Forster and Luschnig, 2012; Luschnig et al., 2006; Nelson et al., 2012; Wang et al., 2006). The transcriptional coactivator Yorkie (Yki) has also been found to be required for proper tracheal tube growth (Robbins et al., 2014), but how a gene, which is mostly implicated in cell proliferation controls growth in a non-proliferating tissue has not been understood.

Yki is a major downstream target of the evolutionarily conserved Hippo signaling pathway, and controls organ size by suppressing proliferation and promoting apoptosis (Halder and Johnson, 2011). Active Hippo signaling phosphorylates Yki, inhibiting its translocation into the nucleus and its transcriptional activity. Upon inactivation of the kinase cascade, non-phosphorylated Yki enters the nucleus and promotes growth by transcriptional activation of genes that inhibit apoptosis, e.g. *Death-associated inhibitor of apoptosis 1* (*Diap1*), or those stimulating proliferation, e.g. Myc.

Since cell proliferation genes are transcriptional targets of Yki and its conserved vertebrate ortholog YAP (Yes-associated protein), Yki/YAP activity has to be tightly regulated, which is largely mediated by controlling its subcellular localization (nuclear versus cytoplasmic). Several upstream mechanisms are known that control Yki/YAP localization, including its phosphorylation (as part of the evolutionarily conserved Hippo pathway) (Fulford et al., 2018), its interaction with tight junction protein complexes that stabilize the cytoplasmic Yki/YAP pool (Skouloudaki and Walz, 2012) and regulate Yki/YAP cytoplasm-to-nucleus translocation in proliferating epithelial tissues (Wang et al., 2011; Zhao et al., 2011), or cellular and extracellular mechanical forces (Totaro et al., 2018).

Here we describe a novel mechanism to control Yorkie cellular localization in a non-proliferating tissue, the tracheal system of *Drosophila*. We discovered that Yki, which has been shown to control tube length expansion (Robbins et al., 2014) does so by interacting with Twinstar (Tsr), the *Drosophila* orthologue of vertebrate Cofilin. Tsr keeps Yki apically in tracheal epithelial cells and thereby prevents its nuclear localization. In addition we show that *yki* is required for the generation of water impermeable tubes. This function is controlled by the Yki-target gene *Alas*, which encodes an enzyme that regulates the structure of the apical extracellular matrix (aECM) (Shaik et al., 2012). Thus, our data provide mechanistic evidence how Yki controls tube elongation and impermeability in the tracheae.

## Results

### Yorkie controls tracheal tube length and proper gas-filling

*Drosophila* embryos, bearing a deletion of the whole *yki* locus (Huang et al., 2005) (*yki*^*B5*^ loss-of function allele, henceforth referred to as *yki*), die at late embryonic stages with elongated tracheal tubes (Fig. 1, A and B) (Huang et al., 2005; Robbins et al., 2014). Staining for chitin confirmed that the length of the major tracheal tubes, the dorsal trunks (DTs), of late stage 16 *yki* mutant embryos is significantly increased compared to wild type (Fig. 1 B). We noticed that beside extended length the DTs fail to clear luminal liquid and do not fill with gas (Fig. 1, C and D). Both phenotypes, i.e. increased tube length and defects in gas filling, were suppressed upon tracheal-specific expression of a cDNA, which encodes a Yki protein C-terminally tagged with V5 (Yki-V5) (Fig. 1, A-D). However, and in contrast to published data (Robbins et al., 2014), we only observed partial rescue of the tube length phenotype of *yki* mutants upon tracheal expression of *Diap1* [*Death-associated inhibitor of apoptosis 1*; also known as *thread* (*thr*)], a well-established Yki-target gene (Fig. 1 E). These results indicate that *yki* is required to limit tracheal tube length and to promote gas-filling in *Drosophila* embryos.

**Figure 1.**
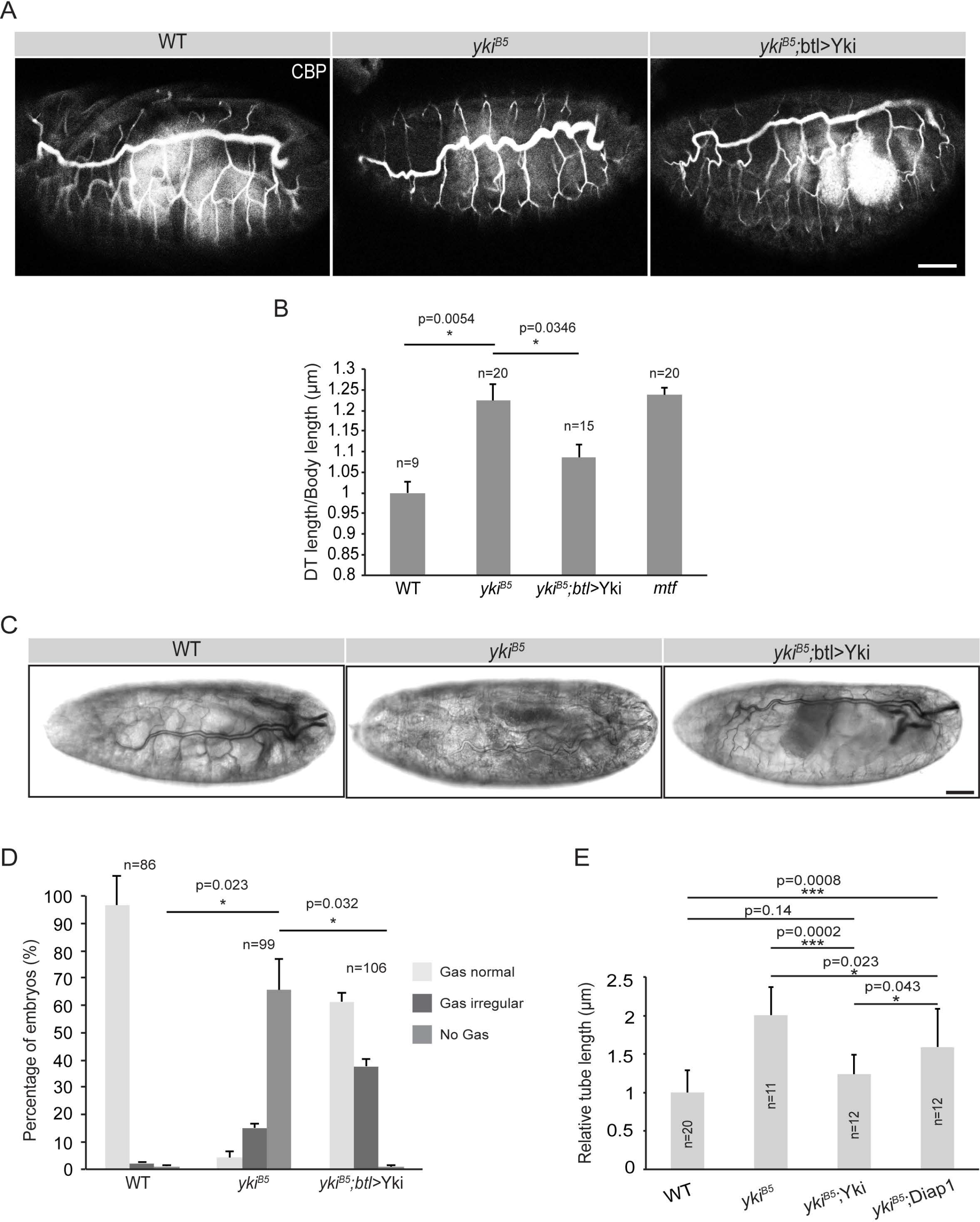
Yki is required to restrict tracheal tube length. (**A**) Wild type (WT), *yki*^*B5*^, *yki*^*B5*^;*btl*>Yki late stage 16 embryos, stained with the chitin binding probe (CBP). Note the convoluted dorsal trunk (DT) tubes in *yki*^*B5*^ mutant embryos. Tracheae-specific expression of *yki* rescues this phenotype. Scale bar: 50 μm. (**B**) DT length is significantly longer in *yki*^*B5*^ mutants compared to that of wild type (WT) embryos or *yki*^*B5*^ mutant embryos expressing Yki under the control of the tracheal driver *btl*-GAL4. Tube length is expressed as a ratio of DT length (metamere 6-10) to body length, normalized against wild type embryos (ratio taken as 1). Included are the measurements of tube length of *mtf* mutants, a gene encoding a bona fide septate junction component. (**C**) Wild type (WT), *yki*^*B5*^, *yki^B5^;btl>*Yki late stage 17 embryos. Gas-filling of the DT lumen is observed in wild type and *yki^B5^;btl>*Yki, but not in *yki*^*B5*^ mutant embryos. Scale bar: 20 μm. (**D**) Plot showing the percentage of *yki*^*B5*^ mutant embryos with gas filling defects, which is significantly different from wild type (WT) and *yki^B5^;btl>*Yki embryos. (**E**) The relative tube length of *yki*^*B5*^ mutant embryos is rescued to different extents by expression of either *yki* or *diap1* by *btl*-GAL4.

### Yorkie is required for formation of impermeable tubes

Liquid impermeability of tracheal tubes has been shown to be controlled by septate junctions (SJ). Loss of SJ components result in leakiness of the tube and impaired gas filling. However, in agreement with previous reports (Robbins et al., 2014), canonical SJ proteins, such as Yurt, Coracle, Megatrachea, Varicose, Fasciclin III and Contactin appeared to be localized correctly to SJs in *yki* mutant tracheae (Fig. 2, A-D and Fig. S1). In agreement with this observation, electron microscopic analysis revealed that the structure of the SJs is not affected in *yki* mutant tracheae and showed proper organization of the ladder-like septa similar to wild type embryos (Fig. 2, A, B, D). This phenotype was rescued by the tracheal expression of a *yki* cDNA (Fig. 2, C, D). Yet, in contrast to recently published data (Robbins et al., 2014), we observed that in approximately 80% of *yki* mutant embryos injected with 10 kDa Dextran to test for the barrier function of SJs, the dye diffused into the lumen of the DTs, which never occurs in wild-type embryos (Fig. 3, A, B and D). Stage 17 embryos expressing a *yki* transgene fail to fill the tracheal lumen with Dextran (Fig. 3, C, D). This indicates that the tracheal epithelium in *yki* mutants is not tightly sealed and that this mechanism is *yki*-dependent. The dye diffused gradually into the tracheal lumen of *yki* mutants, with higher amounts observed 240 min after injection (Fig. 3, E-G” and H). This defect is much milder than that observed in embryos mutant for *melanotransferrin* (*mtf*) (Fig. 3, G-G”), a gene encoding a bona-fide SJ component, in which the dye filled the tracheal lumen already after 20min (compare Fig. 3, F-F”).

**Figure 2.**
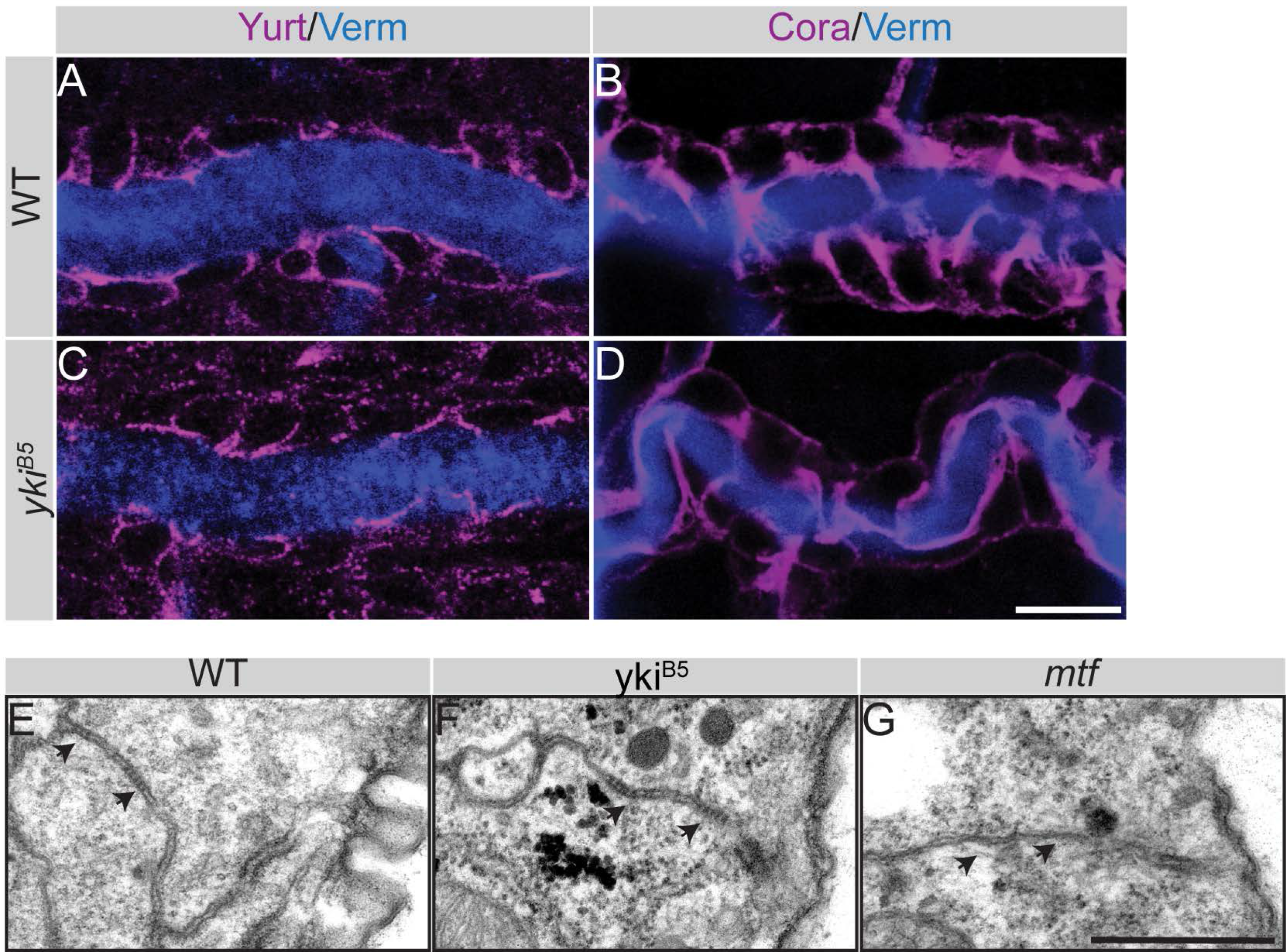
Septate junctions and luminal matrix are not affected in *yki* mutant tracheal tubes. (**A-D**) Embryonic trachea of wild type (WT) and *yki*^*B5*^ mutants of stage 17 stained for the core septate junction components, Yurt and Cora (magenta) and the luminal matrix protein Verm (blue). Yurt (A,C) and Cora (B,D) staining appear normal and comparable to that of wild type (WT) embryos. Scale bar: 10 μm. (**E-G**) Transmission electron microscopy of stage 16 embryonic tracheae of wild type (WT) (E), *yki*^*B5*^ (F) and *mtf* (G) mutants. Electron dense septa (arrows) are comparable in wild type (WT) and *yki*^*B5*^, but invisible in *mtf* mutants. Scale bar: 0.5 μm.

**Figure 3.**
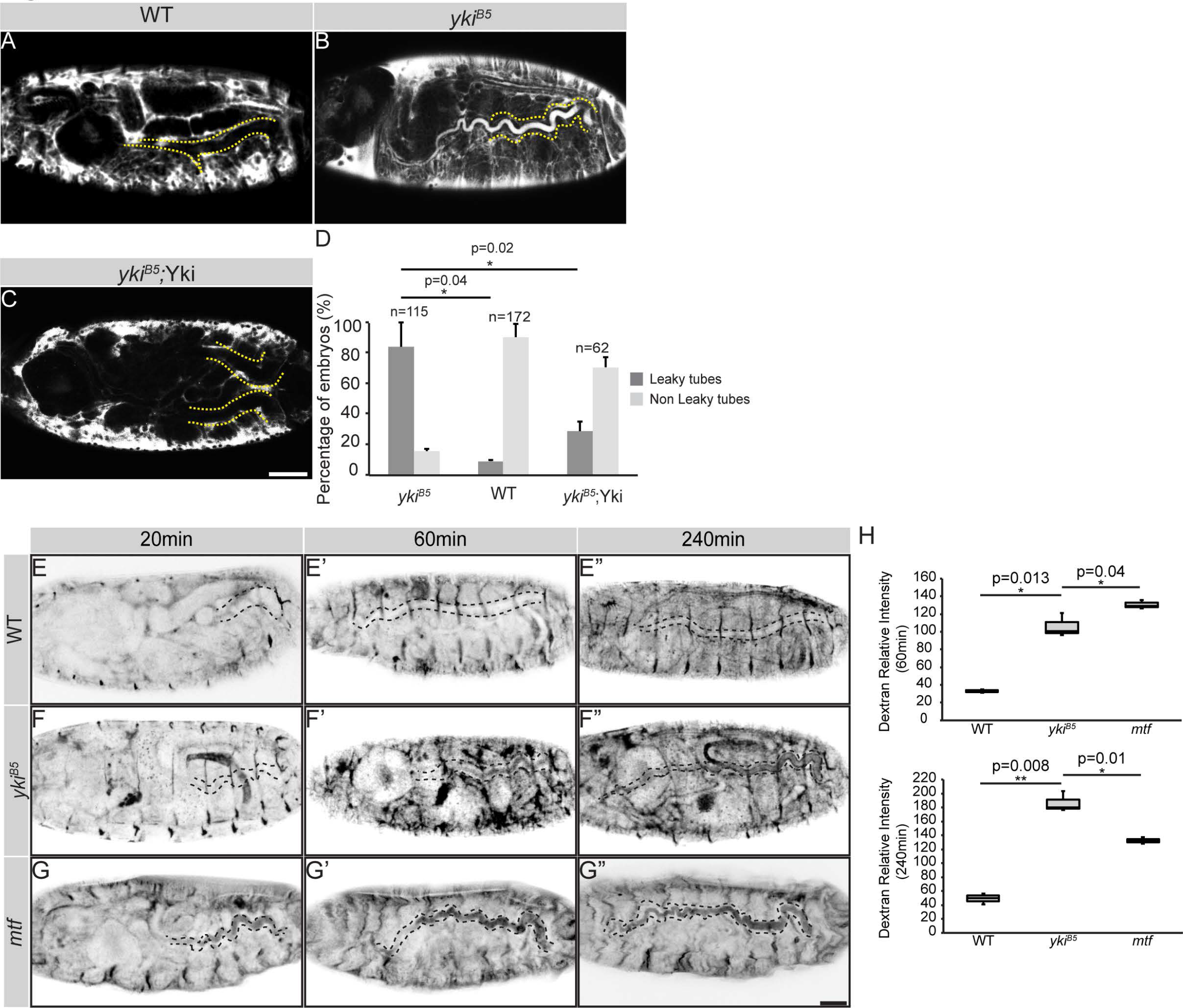
Yki is important for transepithelial barrier function. (**A-C**) Fluorescent 10kDa Dextran injected into the body cavity of embryos at stage 17 does not enter the tracheal lumen (dotted yellow line) of wild type embryos (A). In contrast the dye leaks into the tracheal lumen in *yki*^*B5*^ (B) mutant embryos, indicating a loss of paracellular barrier. In *yki*^*B5*^ embryos expressing Yki in the tracheae, the dye is excluded from the lumen, indicating that *yki* is required in the tracheae to maintain the barrier function (C). Scale bar: 50 μm. (**D**) Plot representing the percentage of *yki*^*B5*^, WT and *yki^B5^;*Yki embryos with leakage defects. (**E-G”**) Time series of Dextran injected embryos. Dextran accumulates gradually in the tracheal lumen of *yki*^*B5*^ mutant embryos as compared to *mtf* mutant embryos, where the dye accumulates a few minutes after injection and stays unchanged over time. Scale bar: 100 μm. (**H**) Quantification of the relative luminal Dextran intensity 60 min (upper plot) and 240 min (lower plot) after injection.

The water-tightness of tracheal tubes is not only ensured by SJs, but also depends on the proper deposition of a chitinous apical extracellular matrix (aECM). Since SJs did not appear impaired in *yki* mutants, we asked whether loss of *yki* affects the secretion of the chitin deacetylases, Serpentine (Serp) and Vermiform (Verm). However, the luminal deposition of Serp and Verm is not perturbed in *yki* mutants (Fig. 2, A-D and Fig. S2, A-B”), pointing to a novel, SJ and luminal matrix deposition-independent function of Yki in the control of tracheal tube impermeability.

### Yorkie is required for proper cross-linking of tracheal apical ECM proteins

Beside proper SJs and secretion of the chitin-modifying enzymes Serp and Verm, formation of an intact apical chitin lattice is necessary to ensure impermeability of tracheal tubes (Shaik et al., 2012). Therefore, we examined the aECM in *yki* mutant tracheae in more detail. The insect cuticle is an apical extracellular waterproof barrier, deposited by epithelial organs such as the epidermis, the tracheae, the fore- and hindgut, which protects against dehydration and infection and plays a crucial role in organ morphogenesis (Moussian, 2010; Ozturk-Colak et al., 2016). The cuticle is composed of three layers, the outermost envelope, followed by the epicuticle and the procuticle. In the tracheae, the epi- and procuticle form the so-called taenidia, chitinous ridges lining the lumen. These layers are formed by proteins, chitin and lipids. Some of the proteins are cross-linked in order to form an impermeable layer. Three lines of evidence suggested that an impaired function of the δ-aminolevulinate synthase gene *Alas* (Shaik et al., 2012) is responsible for fluid leakage and gas filling defects in *yki* mutants. First, ChiP-seq data from Nagaraj *et al*. showed that Yki binds the regulatory region of the *Alas* gene in *Drosophila* imaginal discs (Nagaraj et al., 2012). Second, similarly to *yki*, *Alas* is required for air-filling of the tracheal tubes and for the maintenance of tracheal impermeability at later stages of embryogenesis (Shaik et al., 2012). Third, barrier defects observed in *Alas* mutants occur despite structurally and functionally normal SJs (Shaik et al., 2012), similar to *yki* mutants. This prompted us to investigate a possible functional relationship between *yki* and *Alas*. *Alas* mutant tubes display reduced amount of cuticular di-tyrosine bonds, resulting in loss of cuticular impermeability and defects in tracheal air-filling (Fig. 4, A, A’, E and G) (Shaik et al., 2012). These phenotypes were strongly suppressed upon tracheal-specific expression of an *Alas* cDNA (Fig. 4, B, B’, E, F and G). Di-tyrosine bonds were also reduced in amount in *yki* mutants (Fig. 4, C, C’, E and G), suggesting that *Alas* may act downstream of *yki*. In fact, gas-filling and trans-epithelial barrier defects were rescued by expression of *Alas* in tracheal cells of *yki* mutants (Fig. 4, D, D’, E, F and G), while normal tube length was not restored (data not shown). Furthermore, *Alas* mRNA is almost 2-fold decreased in *yki* mutants (Fig. 4 H), corroborating the notion that Yki is a transcriptional activator of *Alas*. Interestingly, *duox* mRNA, which encodes an enzyme that catalyzes the oxidation of two tyrosines to di-tyrosine (Edens et al., 2001) and which is also expressed in developing tracheae (Yao et al., 2017) did not show any significant difference in *yki* mutants (Fig. 4 H). This led us to suggest that Yki regulates the di-tyrosine network through *Alas*. To test whether Yki is a regulator of di-tyrosine bridges between extracellular proteins in other cuticular organs, we used *ptc*-Gal4 to express Yki-V5 (Fig. S3, A -A”) or knock it down by *yki* RNAi (Fig. S3, B–B”) along the anterior-posterior (AP) boundary of wing imaginal discs. In Yki-expressing discs, the level of di-tyrosine was considerably increased (Fig. S3 A’ arrows, D), whereas di-tyrosine was reduced upon *yki* knock-down (Fig. S3 B’ arrows, D), as compared to the control (Fig. S3, C-C”).

**Figure 4.**
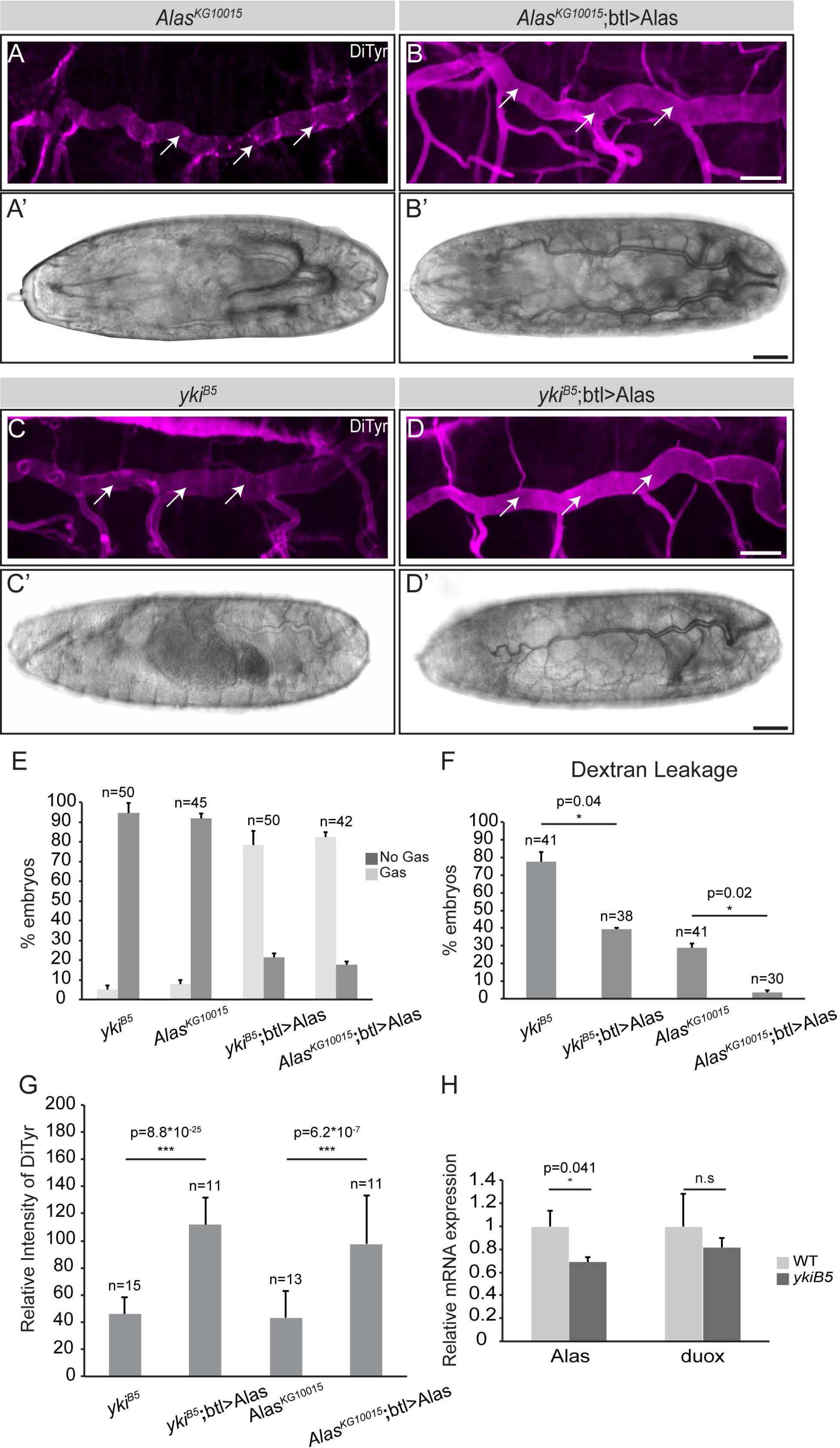
The apical barrier breaks down in *yki* mutant embryos. (**A-D**) Projections of confocal sections of tracheal dorsal trunks of stage 17 embryos. The di-tyrosine network marking the apical barrier (magenta) is reduced in *Alas*^*KG10015*^ (A) and *yki*^*B5*^ (C) mutant embryos, whereas the signal is markedly increased in mutant embryos expressing Alas by *btl*-GAL4 (B, D). White arrows indicate DiTyr apical staining. Scale bar: 20μm. (**A’-D’**) Whole mount embryos at early stage 17. The DT of *Alas*^*KG10015*^ (A’) and *yki*^*B5*^ (C’) mutant embryos are not air-filled. In contrast, expression of Alas with the tracheal-specific driver *btl*-GAL4 rescues the air filling defects of both mutants (B’, D’). Scale bar: 20μm. (**E**) Plot showing the percentage of *Alas*^*KG10015*^ and *yki*^*B5*^ mutant embryos with gas-filling defects, and significant rescue of this defect in both mutants upon tracheal-specific expression of Alas. (**F**) Plot showing the percentage of embryos with defects in the barrier function of the DT, as measured by 10kDa-Rhodamine-dextran leakage into to lumen. (**G**) Quantification of anti-Dityrosine intensity as a measure for the apical extracellular barrier. (**H**) Quantitative real-time RT-PCR showing a significant difference in *Alas* mRNA levels between wild type (WT) and *yki*^*B5*^ mutants at stage 17. No significant difference was detected in *duox* mRNA levels.

To summarize, we provide evidence that Yki acts through transcriptional activation of *Alas* to regulate extracellular di-tyrosine-dependent barrier formation in the tracheae, required for forming waterproof tracheal tubes.

### Yorkie controls tracheal tube length by regulating the actin-depolymerizing factor Twinstar/Cofilin

In contrast to *yki* mutants, *Alas* mutants do not exhibit over-elongated DTs (Fig. 4, A, C and Fig. S4, A-C), suggesting that Yki-mediated regulation of tube expansion is independent of *Alas*. Additionally, the only partial rescue of the tube length in *yki* mutants upon *Diap1* expression points to an additional, *Diap1*-independent function of Yki in determining tube size. Several genes are known to restrict tracheal tube length (Hayashi and Kondo, 2018) including those encoding SJ proteins (Zuo et al., 2013), chitin-modifying enzymes (Luschnig et al., 2006; Wang et al., 2006), polarity proteins regulating apical membrane growth (Dong et al., 2014; Laprise et al., 2006) and cytoskeleton proteins and their regulators [recently reviewed in (Hayashi and Kondo, 2018; Ochoa-Espinosa et al., 2012)]. As shown above, *yki* mutant tracheae bear normal SJs and secrete Verm and Serp normally into the lumen. Localization of the apical determinant Crb and aPKC were apparently not affected in *yki* mutants (Fig. S5, A-B and A’-B’). Finally, genetic interaction experiments between *yki* and *Src42a*, which encodes a tyrosine kinase, revealed that the two genes act in parallel pathways to regulate tracheal tube length (Robbins et al., 2014). Taken together, our analysis identified a function of Yki in the control of tracheal tube length, independent of SJs, luminal matrix deposition and apical polarity.

To unveil the molecular mechanism by which Yki regulates tube length expansion, we immunoprecipitated Yki-V5, expressed in tracheae, and performed mass spectrometry analysis. One of the proteins found to associate with Yki was Twinstar (Tsr), the *Drosophila* ortholog of vertebrate Cofilin (Table S6 A). Recent studies have implicated Tsr in cell survival, tissue growth and tissue integrity of *Drosophila* wing imaginal discs by regulating Yki and JNK signaling (Ko et al., 2016). To further confirm that Yki and Tsr interact physically, we examined their interaction using several assays in embryonic tracheae and wing imaginal discs. Overexpressed Yki-V5 co-precipitated Tsr from protein lysates of embryonic tracheae (Fig. 5 A). Further, in-situ proximity ligation assay (PLA) (Soderberg et al., 2006) (Fig. 5 B-C”) and Bimolecular fluorescence complementation (BiFC) (Hu et al., 2002) (Fig. 5 D) both confirmed that the two proteins, when expressed in wing imaginal discs, were found in close proximity. From these results we concluded that Yki and Tsr physically interact to form a protein complex.

**Figure 5.**
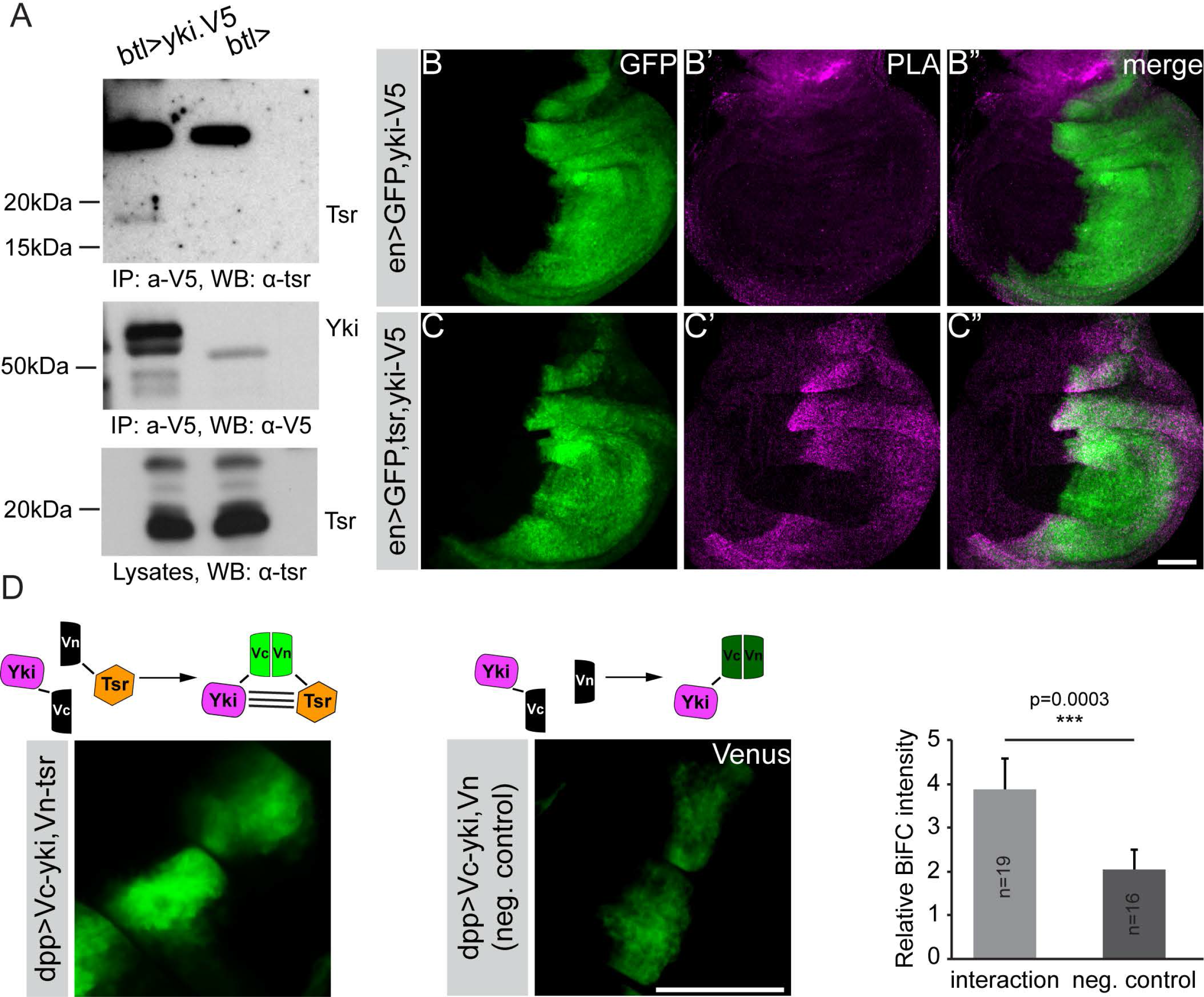
Tsr and Yki interact. (**A**) Tsr co-immunoprecipitates with Yki from embryo lysates expressing Yki-V5 in tracheal cells.*btl*-GAL4 alone was used as a negative control. IP, immunoprecipitate; WB, western blot. (**B-C”**) Proximity ligation assay of Tsr and Yki in wing imaginal discs. Wing discs from *en*>GFP,Yki-V5 (control) and *en*>GFP, Yki-V5, Tsr were labeled with anti-V5 and anti-Tsr to perfom PLA assays. The Yki-V5 expression domain is marked by GFP. (**D**) Bimolecular Fluorescence Complementation of Yki and Tsr complexes in wing imaginal discs. The diagram depicts the relative BiFC intensity of complexes as compared to control. Scale bars: 50μm.

To find out whether Tsr and Yki function together to regulate tube size, we analyzed the function of Tsr in the tracheae, using two different alleles, *tsr*^*k05633*^ and *tsr*^*N96A*^ (Ng and Luo, 2004; Wahlstrom et al., 2001). Strikingly, *tsr* mutant embryos exhibited over-elongated tracheal tubes, similar to *yki* mutants (Fig. 6 A-C, E and Fig. S7 A-C). This phenotype is due to a specific function of Tsr in the trachea, since tracheal-specific expression of a *tsr* cDNA by *btl*-GAL4 rescued the over-elongated tubes of *tsr* mutants (Fig. 6 F-H). In contrast to *yki*, however, *tsr* mutants showed proper tracheal gas-filling (Fig. S7 D and E) and displayed normal paracellular barrier of the tracheal tubes (Fig. S7 F and G). These results indicate that *tsr* functions in tube size control, but not in gas filling.

**Figure 6.**
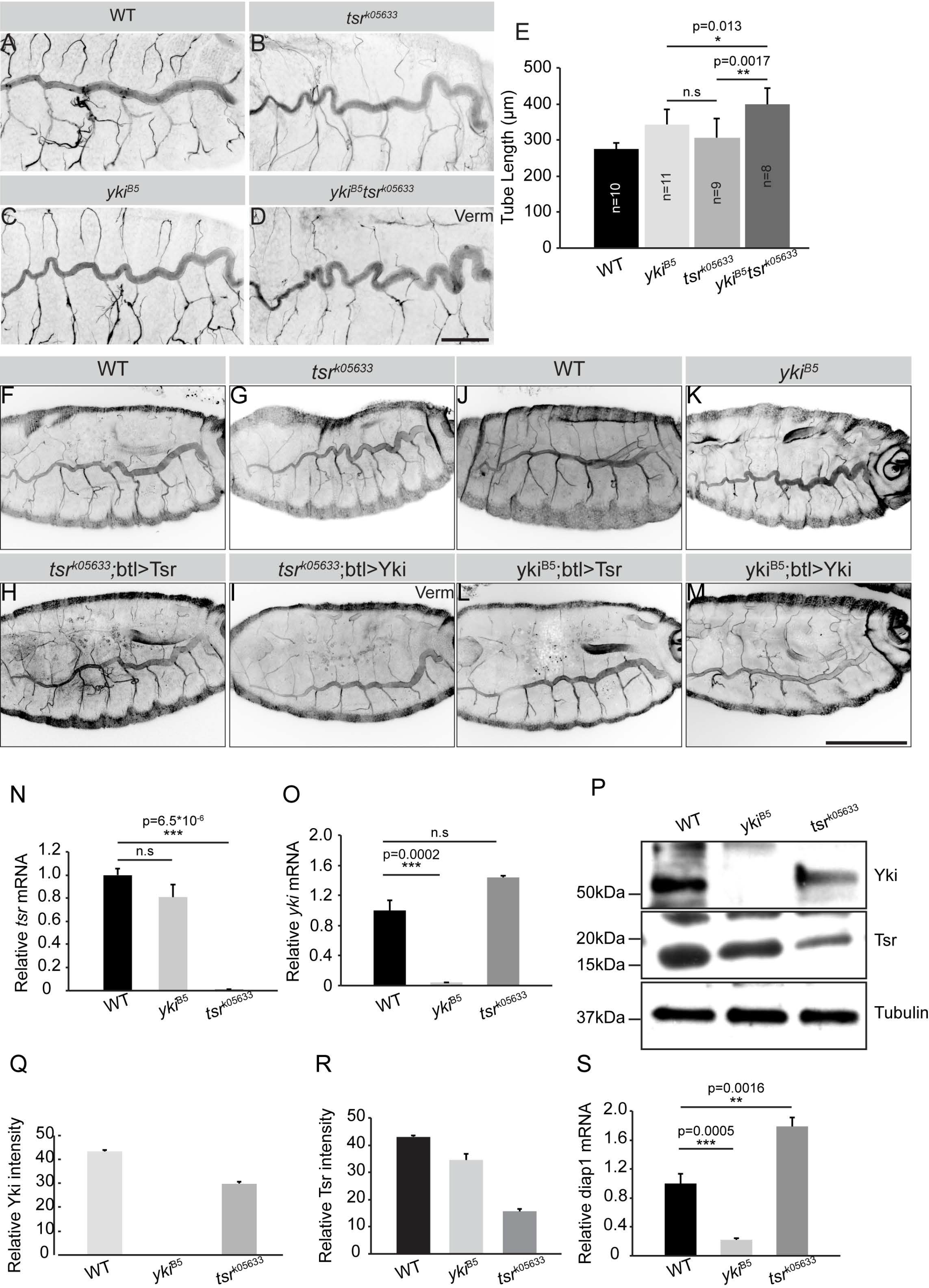
Tsr and Yki act in interconnected pathways to regulate tracheal tube elongation. (**A-E**) Loss-of function of *tsr* (B) causes convoluted DT, similar as loss of *yki* (C). This phenotype is enhanced in *tsr*^*k05633*^; *yki*^*B5*^ double mutants (D). (E) Quantification of DT length of wild type (WT), *yki*^*B5*^, *tsr*^*k05633*^ and *tsr*^*k05633*^; *yki*^*B5*^ mutants. The DT of *tsr*^*k05633*^; *yki*^*B5*^ double mutants is significantly longer than that of *yki*^*B5*^ and *tsr*^*k05633*^ single mutants. Scale bar: 50μm. (**F-M**) Tracheal expression of either Tsr (H, L) or Yki (I, M) using *btl*-GAL4 rescues DT elongation defects of *yki*^*B5*^ and *tsr*^*k05633*^ mutants. Scale bar: 20 μm. (**N**) Relative expression of *tsr* mRNA in wild type (WT), *yki*^*B5*^ and *tsr*^*k05633*^ mutant embryos of stage 17. *tsr* mRNA levels are not significantly altered in the absence of *yki*. (**O**) Relative expression of *yki* mRNA in wild type (WT), *yki*^*B5*^ and *tsr*^*k05633*^ mutant embryos of stage 17. *yki* mRNA levels are not significantly altered in the absence of *Tsr*. (**P**) Western blot of protein lysates from wild type (WT), *yki*^*B5*^ and *tsr*^*k05633*^ mutant embryos of stage 17. Note that the protein levels of Tsr and Yki are reduced in the respective other mutant. (**Q, R**) Quantification of the immunoblot in (P) using Fiji, based on the intensity of Yki (Q) and Tsr (R) protein, normalized to the loading control (alpha-Tubulin) (n=2). (**S**) Relative expression of *diap1* mRNA in wild type (WT), *yki*^*B5*^ and *tsr*^*k05633*^ mutant embryos of stage 17. Results were normalized to an endogenous control (actin-5C). Note that *diap1* is significantly downregulated in *yki* mutants, but significantly upregulated in *tsr* mutants

The similar tracheal phenotypes of *tsr* and *yki* mutants raised the possibility that Yki and Tsr act in the same pathway to control DT length. Strikingly, *yki*, *tsr* double mutants had even longer tubes, as compared to those of *yki* and *tsr* single mutants, suggesting that *yki and tsr* act in parallel pathways to regulate tube size (Fig. 6 D and E). To determine whether there is any connection between the two pathways to regulate tracheal tube length, we expressed a *yki* cDNA in the tracheae of *tsr* mutant embryos. In these embryos tracheal tube length was restored and comparable to that of control embryos (Fig. 6 I). Likewise, expression of a *tsr* cDNA reversed the *yki* tube elongation phenotype (Fig. 6 J-M). These data support that *tsr* and *yki* act in parallel, yet interconnected pathways.

### Tsr regulates Yki nuclear activity

To determine the functional relationship between Tsr and Yki in the developing *Drosophila* airways, we first asked how they cooperate to control tube length. Interestingly, Yki binding sites were also identified in the regulatory sequences of *tsr* (Nagaraj et al., 2012), suggesting a transcriptional regulation of *tsr* by Yki in imaginal discs. However, we did not find any significant alteration in *tsr* mRNA levels between control and stage 17 *yki* mutants by quantitative real-time PCR (Fig. 6 N), while *yki* mRNA was undetectable in *yki* mutants at this stage (Fig. 6 O). Therefore, we asked whether our inability to detect changes in *tsr* mRNA levels are owed to maternal Yki protein, which can be detected in unfertilized eggs (Fig. S8 A). We undertook a biochemical approach and observed a Yki life time of 11-16 hours in S2 cells (half-life of t_1/2_ ≈ 3 h). Assuming the same life time in embryos suggests the absence of Yki protein (maternal and zygotic) in stage 17 *yki* mutants, at which *tsr* transcripts were analyzed (Fig. S8, B-C and Fig. 6, P-R). From this we conclude that *tsr* is not a transcriptional target of *yki* in the embryo (or in the tracheae).

How is then Tsr regulated? Interestingly, in contrast to *tsr* transcripts, the protein levels were reduced in stage 17 *yki* mutant embryos. At the same time, we observed a mutual regulation between Yki and Tsr, in that Yki protein is also reduced in *tsr* mutant embryos (Fig. 6 P-R). To determine whether the Tsr-dependent changes in Yki levels go along with changes in Yki transcriptional activity, we analyzed *in vivo* the transcription of one if its target genes, *Diap1*. *Diap1* mRNA levels were significantly reduced in *yki* mutant embryos but increased about two-fold in *tsr*^*k05633*^ mutants (Fig. 6 S). To determine whether this increase in *diap1* transcription is also observed in the tracheae, we used the *yki* transcriptional reporter, *Diap1-lacZ*-NLS (Ryoo et al., 2002). In fact, the reporter showed stronger expression in tracheal cells of *tsr* mutant embryos (Fig. 7 A and B).

**Figure 7.**
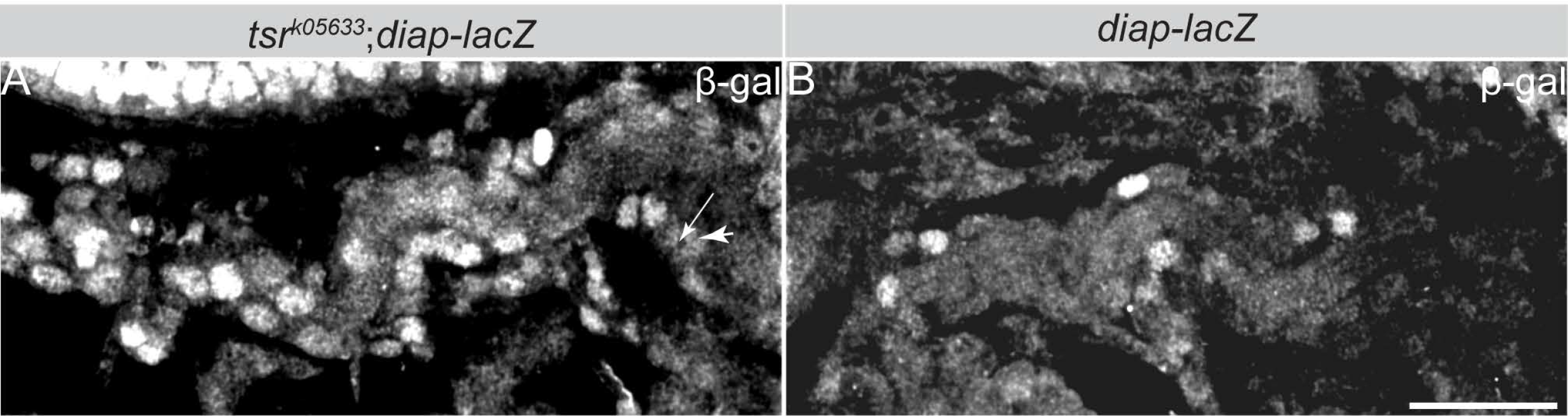
Tsr regulates Yki nuclear localization and the expression of its target gene, *diap1*. (**A,B**) *Tsr* mutant tracheal cells of stage 17 embryos show increased levels of Yki-target gene *diap1-lacZ* (A) as compared to the control (B).

Yki activity is mostly controlled by phosphorylation of S168 (in Yki) and S127/S381 (in Yki/YAP). Phosphorylated Yki/YAP is retained in the cytoplasm, while non-phosphorylated Yki/YAP can enter the nucleus and activate transcription of its target genes (Oh and Irvine, 2011). Using the phosphate-binding tag method (Kinoshita et al., 2006) to capture the phosphorylated form of Yki in embryonic protein lysates, we could not draw any meaningful conclusion due to low p-Yki levels. However, since this regulation seems to be conserved in vertebrates, we addressed this question in HEK293T cells. In this experiment, we used *cofilin* siRNA to downregulate its expression levels. In fact, we observed a reduction in total YAP as well as in p-YAP (indicated by pS127-and pS381-YAP) levels (Fig. S9). These results suggest that both the amount and the phosphorylation status of Yki depend on Tsr.

Taken together, our data suggest that Tsr is a negative regulator of Yki nuclear localization and its transcriptional activity.

### Yorkie and Twinstar cooperate to coordinate apical membrane expansion

Increased tube elongation in *yki* mutants is not caused by an increase in cell number (Fig. S10, A and B), suggesting a different cellular/molecular mechanism, e.g. changes in cell shape. To quantify cell shape, we examined the apical and basal domains of tracheal cells, marked by Uninflatable (Uif) and Perlecan, respectively. In wild-type embryos, the two membrane markers are lined approximately parallel along the tube length (Fig. 8 A-A” and C-C”). In contrast, the DTs of *yki* (Fig. 8 B-B”) and *tsr*^*k05633*^ (Fig. 8 D-D”) mutants showed increased apical membrane with irregular shape, but unaffected basal membrane size, suggesting that only the apical membranes of tracheal cells are over-elongated. Quantification using the adherens junction (AJ) protein Echinoid (Ed) revealed significant increase in the surface of the apical membrane of about 17.2% in *yki* and 18.3% in *tsr*^*k05633*^ mutants (Fig. 8 F, J, M and N), as compared to wild type cells (Fig. 8, E, I, M and N). In line with the mutual stabilization of Tsr and Yki at the protein level (Fig. 6, P-R), tracheae-specific expression of *tsr* or *yki* rescued the *yki* and *tsr*^*k05633*^ mutant expansion defects, respectively (Fig. 8, G, K, M and N). Based on these experiments, we conclude that increase apical membrane size responsible for changes of tracheal epithelial cells that induce changes in tube size through Tsr/Yki.

**Figure 8.**
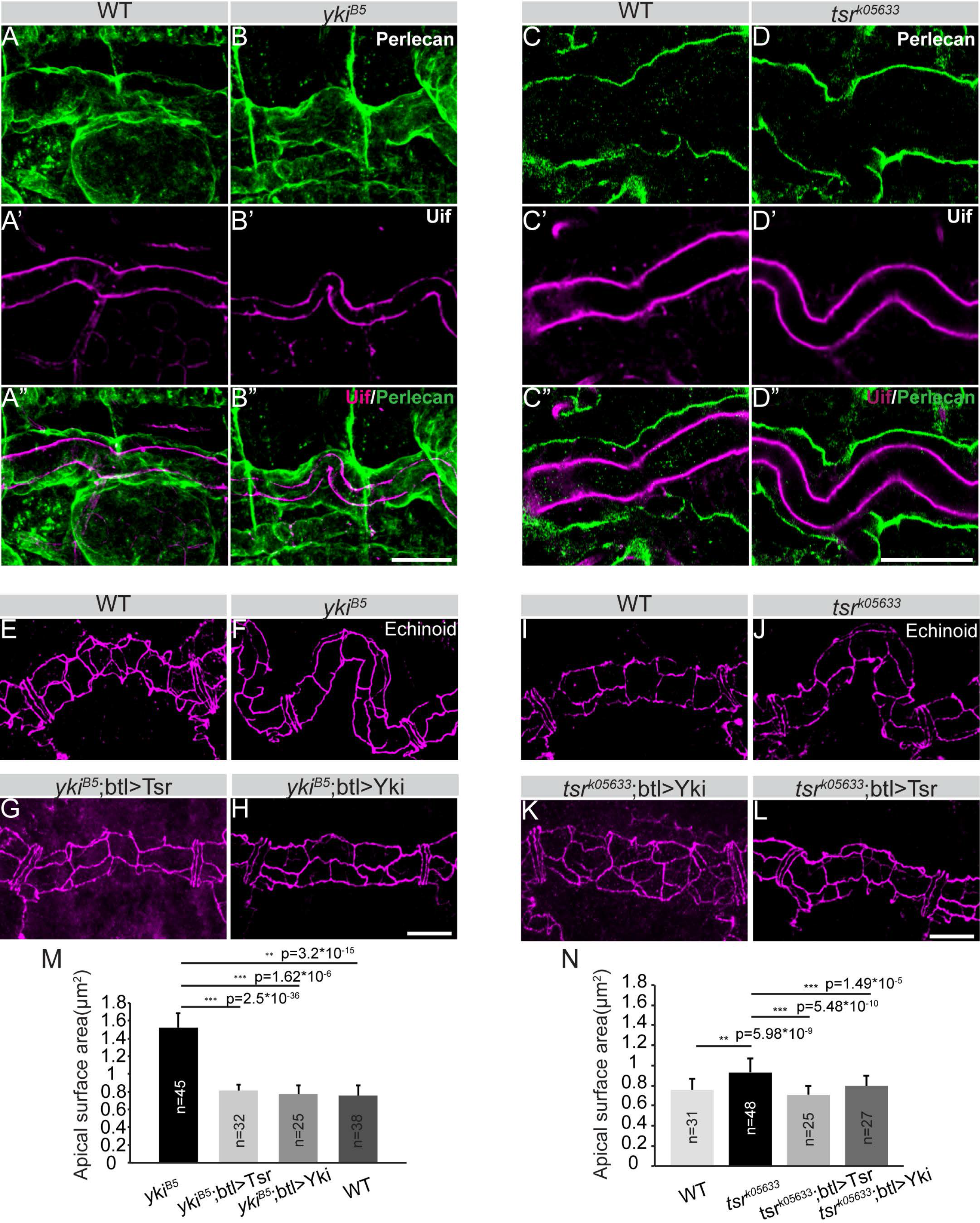
Apical membrane expansion contributes to tube over-elongation in *yki* and *tsr* mutant embryos. (**A-D”**) Stage 17 wild type (WT) (A-A” and C-C”) and *yki*^*B5*^ mutant (B-B” and D-D”) embryos stained with Uif to label the apical membrane (A’-D’ and A”-D” magenta) and Perlecan to label the basement membrane (A-D and A”-D”; green). Scale bars: 20 μm. (**E-L**) Stage 17 embryos stained with Echinoid to outline the apical surface of DT cells. Scale bars: 10μm. (**M**) Quantification of the apical surface of *yki*^*B5*^ and control embryos. A significant increase of the apical surface area is observed in *yki*^*B5*^ mutants compared to wild type (WT). Apical surface area is restored upon tracheal-specific expression of Tsr or Yki in *yki*^*B5*^ mutant embryos. (**N**) Quantification of the apical surface of *tsr*^*k05633*^ and control embryos. A significant increase of the apical surface area is observed in *tsr*^*k05633*^ mutants compared to wild type (WT) embryos. Apical surface area is restored upon tracheal-specific expression of Tsr or Yki in *tsr*^*k05633*^ mutant embryos.

### Loss of Twinstar and Yorkie affect tube length through changes in F-actin organization

We next sought to identify players involved in apical membrane mediated changes in tube size. Loss of *tsr* or *yki* (this study) and increased F-actin polymerization (Sansores-Garcia et al., 2011) both cause cell polarity-independent overgrowth. In addition, increased F-actin at the apical surface enhances Yki-mediated gene expression in wing imaginal discs (Fernandez et al., 2011). Moreover, decreased cortical F-actin can lead to increased expansion of the apical cell membrane due to lack of the formation of a contractile network (Forest et al., 2018; Haigo et al., 2003; Kinoshita et al., 2008; Lee and Harland, 2007; Spencer et al., 2015; Tsoumpekos et al., 2018). These observations suggested a link between the loss of *yki* and F-actin modulation. Tsr is an actin depolymerization factor that catalyzes F-actin disassembly (Bamburg, 1999). Therefore, we set out to investigate actin distribution in *yki* and *tsr*^*k05633*^ mutants. In stage 17 wild type embryos, F-actin (marked by Utrophin and Lifeact) is enriched at the apical cortex of wild type tracheal cells (Fig. 9A, A’, C, C’). F-actin accumulates even more in the apical cortex of DT cells of *tsr*^*k05633*^ (Fig. 9B, B’) 9 A-B’) and *yki* (Fig. 9, D, D’) mutants. To better follow Yki localization, we generated an endogenous CRISPR knock-in line expressing an N-terminal tagged form of Yki (called mKate2-Yki). mKate-Yki flies are homozygous viable and fertile, indicating that the generated *yki* allele is fully functional, as judged by its ability to rescue a *yki* null mutation in transheterozygote animals (Fig. S11 A and B).

**Figure 9.**
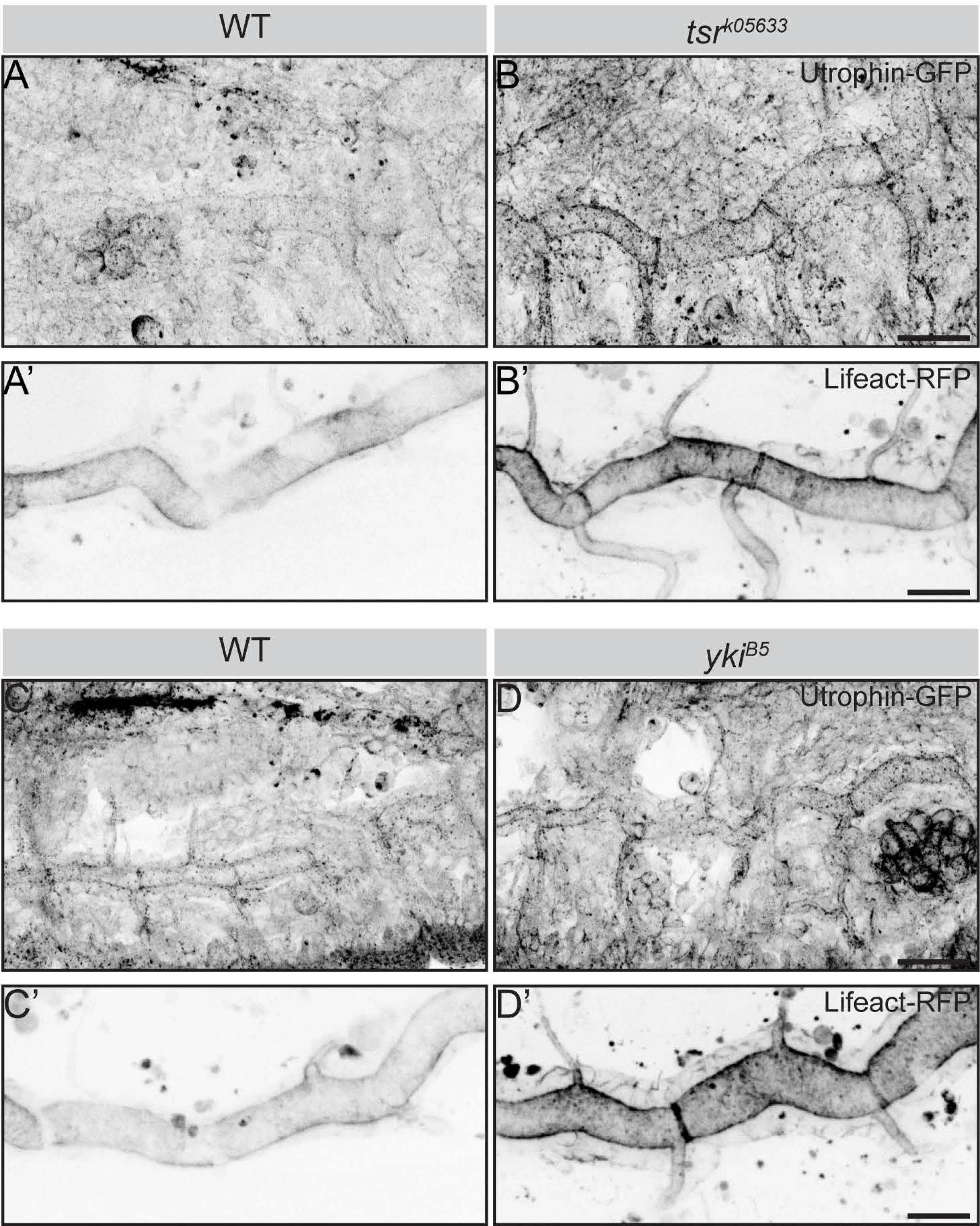
*tsr* and *yki* mutant tracheae exhibit increased apical actin. (**A-D’**) Maximum intensity projections of stage 17 embryos expressing (Utrophin)-GFP (A, B, C, D) and Lifeact-RFP (btl>Lifeact-RFP) (A’, B’, C’, D’) to show the apical F-actin cortex in wild type (WT) (A, A’, C, C’), *tsr*^*k05633*^ (B, B’) and *yki*^*B5*^ (C, C’) mutants. Scale bars: 20 μm.

At 17h AEL (after egg laying), mKate2-Yki starts to accumulate at the apical cortex. This apical enrichment of Yki becomes progressively more prominent at 20h AEL (Fig. 10 A and Fig. S11 C). This pattern mirrors the endogenous Yki localization, since wild type tracheal tubes of stage 17 embryos, stained with a Yki-specific antibody showed a similar apical enrichment, which was hardly detected in *yki* mutant embryos (Fig. 10 B and C). To quantify the amount of Yki in the cortex of tracheal cells in comparison to that in other cellular compartments, such as the nucleus and the residual cytoplasm (Fig. 11 A), we performed Fluorescence Correlation Spectroscopy (FCS) analysis in confocal sections of the cortical cellular domain and in sections of the non-cortical cytoplasm in tracheae of stage 17 embryos (Fig. 11 B). FCS is a well-established method with single-molecule sensitivity to study the dynamic behavior of diffusing fluorescent molecules in live cells or tissues, and extract information about their absolute concentrations and molecular movement (diffusion) within cells or cellular compartments (Vukojevic et al., 2010; Vukojevic et al., 2005). In this way, molecular events, such as binding of protein molecules to larger, immobile structures, resulting in retardation of their diffusion, can be studied with very high sensitivity. A summary on FCS methodology is outlined in Supplement 2. We performed FCS measurements using either a rescuing construct of *yki*, Yki-GFP (Oh and Irvine, 2008) or the endogenously tagged mKate2-Yki. The concentration of Yki-GFP and mKate2-Yki was significantly higher in the apical cortex of wild-type tracheal cells and decreased from the cortex to the cytoplasm to the nucleus (Fig. 11 C, E and G). FCS allowed us to also discern the relative fractions of fast-diffusing *versus* slowly-diffusing Yki-GFP molecules in these compartments, which indicate the percentage of the total Yki molecules that are presumably docked to larger immobile structures, as compared to those which are freely moving in each of the cellular compartments. A higher fraction of slowly diffusing Yki-GFP and mKate2-Yki molecules was observed in the cortex, whereas less Yki-GFP and mKate2-Yki molecules were found in the cytoplasm and the nucleus (Fig. 11 D, F and H). From this, we conclude that in tracheal cells a higher fraction of Yki is stabilized at the apical cortex. Based on our results, Tsr is a likely candidate accounting for this stabilization.

**Figure 10.**
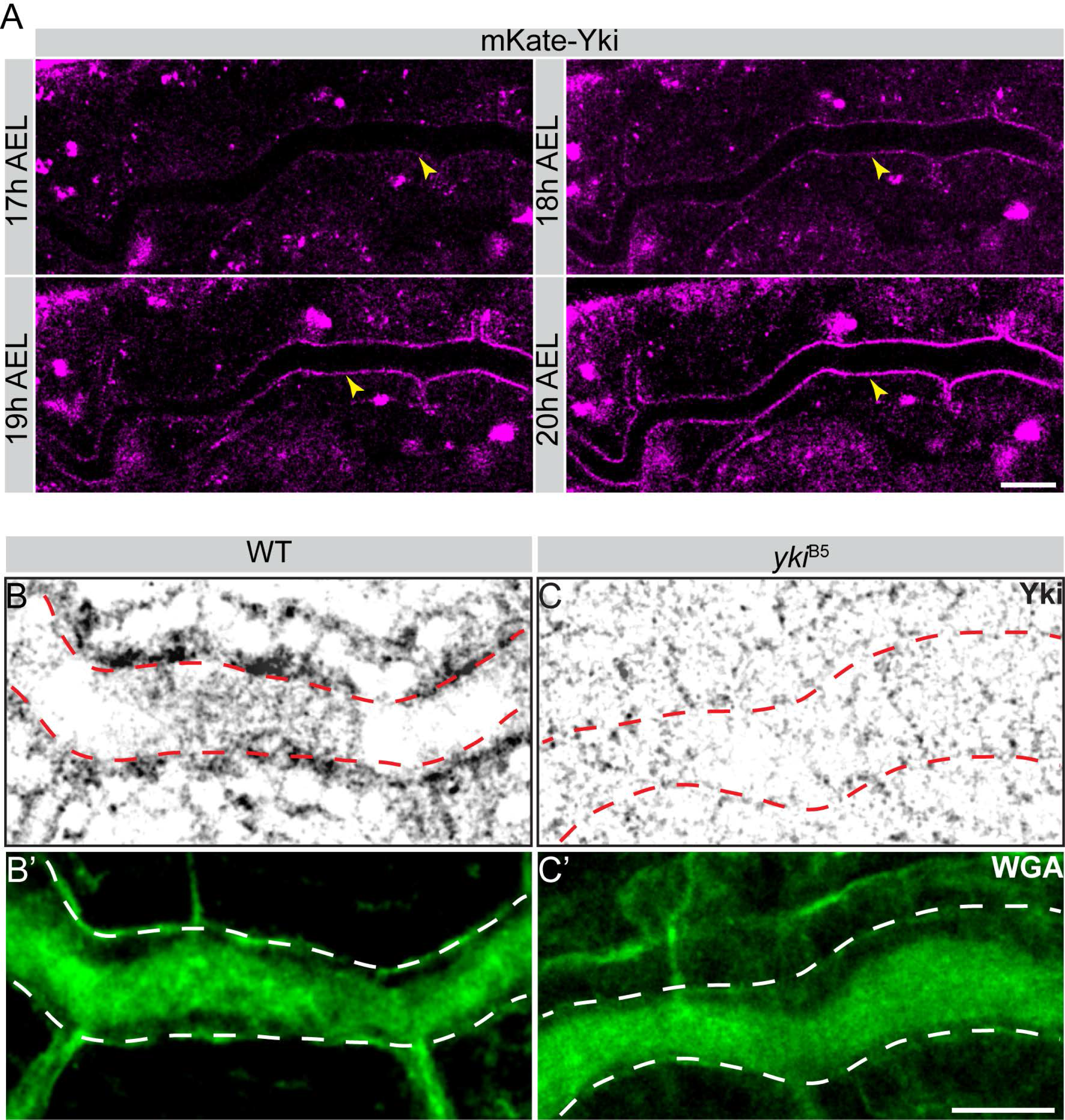
Yki is enriched apically in DT cells. (**A**) Live imaging of Yki dynamics during late tracheal development, using mKate2-Yki. Apical Yki intensity increases with time. Scale bar: 100 μm. (**B-C’**) Confocal images of wild type (WT) (B, B’) and *yki*^*B5*^ mutant (C, C’) embryos stained with anti-Yki (B, C) and WGA (green) to label the lumen (dashed lines outline the lumen) (B’, C’). In wild type (WT) embryos, Yki is enriched at the apical cortex (marked by red dashed line) of tracheal cells (B), whereas *yki*^*B5*^ mutants completely lack Yki cortical labeling (C). (B’, C’). Scale bar: 10 μm.

**Figure 11.**
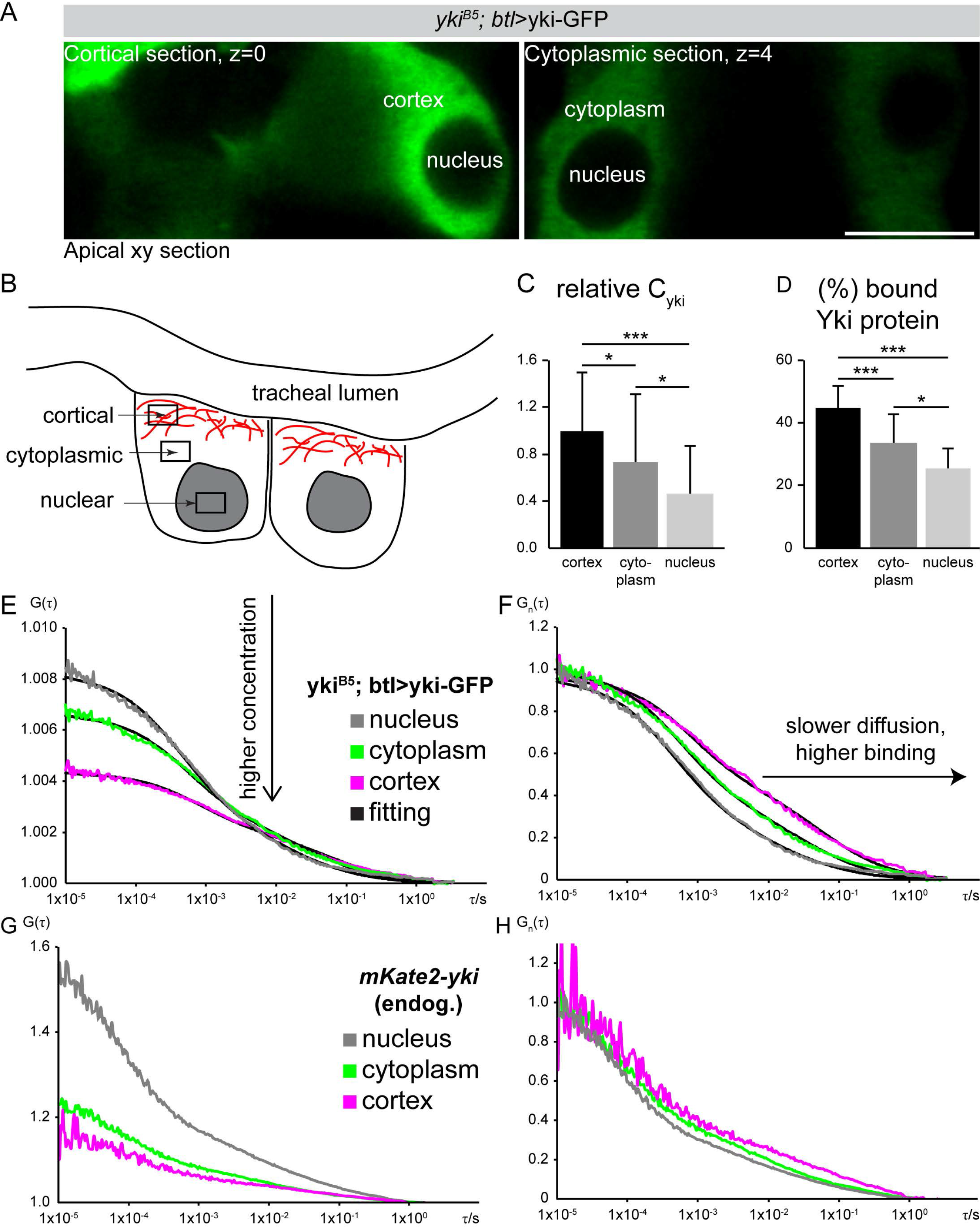
Yki shows the highest concentration at the apical membrane of DT cells. (**A**) Cortical and cytoplasmic sections of the tracheal lumen of stage 17 embryos expressing Yki-GFP by *btl*-Gal4. Scale bar: 5 μm. (**B**) Schematic representation of the different tracheal cell areas, in which the concentration of Yki-GFP was determined by Fluorescence Correlation Spectroscopy (FCS). (**C**) Relative Yki-GFP concentrations, as determined by FCS in the cortical and cytoplasmic sections, as well as in the nucleus. (**D**) Relative fractions of slowly diffusing Yki-GFP molecules, as measured by FCS. A relatively higher amount of slowly-diffusing Yki-GFP molecules is found in the cortex, as compared to the cytoplasmic area or the nucleus, suggesting more pronounced interactions and binding in the cortex. (**E**) Average FCS curves of Yki-GFP in the cortex, cytoplasm and nucleus. The concentration increases from nucleus to cytoplasm to cortex, as shown also in (C). (**F**) Normalized average FCS curves to the same amplitude, *G*(*τ*) = 1, allow comparison of the diffusion of Yki-GFP in the investigated cellular compartments. Yki-GFP displays increasingly slower diffusion from the nucleus to the cytoplasm to the cortex (n=36 cells). (**G**) Average FCS curves of mKate2-*yki* in the cortex, cytoplasm and nucleus. The concentration increases from nucleus to cytoplasm to cortex. (**H**) Normalized average FCS curves to the same amplitude, *G*(*τ*)= 1, allow comparison of the diffusion of mKate2-*yki* in the investigated cellular compartments. mKate2-*yki* displays increasingly slower diffusion from the nucleus to the cytoplasm to the cortex (n= 27 cells).

Taken together, our results indicate a synergy of Tsr and Yki in the regulation of the actin cytoskeleton to modulate the size of the apical surface of tracheal cells and thereby restrict tube elongation.

## Discussion

Data presented in our study reveal a dual function of *Drosophila* Yki in tracheal development. First, Yki is required to ensure cuticle water-tightness and second, to restrict tube length.

We show that *yki* mutant embryos fail to generate functional gas-filled airways, due to improper establishment of the apical extracellular di-tyrosine network, a structural constituent of the apical ECM, important for tissue integrity. Without excluding additional mechanisms, we have attributed this function to *Alas*, mutations of which phenocopy the gas-filling defects of *yki* mutants. We show that *Alas* is a transcriptional target of *yki* and overexpression of *Alas* rescues the apical intercellular barrier abnormalities and gas-filling defects of *yki* mutant embryos, but not the abnormal tube elongation. Therefore, we conclude that *yki* regulates tube impermeability through *Alas*.

Second, results presented here are the first to unveil a molecular mechanism by which Yki restricts tube length in *Drosophila* airways. We identified *Drosophila* Tsr/Cofilin as the most abundant Yki interactor in tracheal tissue. Tsr/Cofilin has been shown to bind to both monomeric globular (G)-actin and filamentous (F)-actin, and severs F-actin, thus causing its depolymerization (Andrianantoandro and Pollard, 2006; Bernstein and Bamburg, 2010; Moon and Drubin, 1995). Therefore, regulation of Tsr/Cofilin is critical for adjusting actin dynamics during tissue morphogenesis (Kiuchi et al., 2011). Here, we provide data to suggest that Tsr forms a protein complex with Yki to restrict tracheal tube elongation through regulation of actin polymerization. Depletion of either *tsr* or *yki* induces increased cortical F-actin. As a consequence, the apical membrane expands, and thus tubes grow in length. This finding is in line with studies showing that disassembly of F-actin fosters apical constriction in cells of the early *Drosophila* embryo (Jodoin et al., 2015). The observed Yki-Tsr interplay is consistent with findings in mammary epithelial cells (MEC) (Aragona et al., 2013), squamous carcinoma cells (SCC) (Kanellos et al., 2015) and in cells of the *Drosophila* wing epithelium (Ko et al., 2016). Cofilin/Tsr-depletion in these cells induced upregulation of YAP/TAZ/Yki target genes, such as mammalian *CTGF* (Connective Tissue Growth Factor) or *Drosophila Diap1* and *expanded* (Aragona et al., 2013; Ko et al., 2016). Similarly, depletion of *capulet*, an inhibitor of actin polymerization, or overexpression of a constitutively active allele of *diaphanous*, encoding a *Drosophila* formin, which nucleates actin, results in increased F-actin bundles and tissue overgrowth (Sansores-Garcia et al., 2011). However, in other cases, decreased cortical F-actin leads to increased expansion of the apical cell membrane due to lack of the formation of a contractile network (Forest et al., 2018; Haigo et al., 2003; Kinoshita et al., 2008; Lee and Harland, 2007; Spencer et al., 2015; Tsoumpekos et al., 2018), pointing to cell type-specific consequences of apical actin modulation.

How does accumulation of F-actin control Yki activity and thus contribute to organ growth? Several lines of evidence support the notion that mechanical forces, transmitted through the ECM, junctions or the cytoskeleton, act upstream as regulators of Yki in both *Drosophila* and human cells during development (Dong and Hayashi, 2015; Elbediwy and Thompson, 2018; Schroeder and Halder, 2012; Sun and Irvine, 2016). For example, spectrins, large cytoskeletal proteins and major constituents of the membrane-associated cytocortex (Bennett and Baines, 2001), repress nuclear Yki/YAP activity and therefore tissue growth (Fletcher et al., 2015; Forest et al., 2018). However, our results suggest that the relation between Yki activity and the actin cytoskeleton is not unidirectional and that Yki feeds back to the cytoskeleton via the regulation of Tsr (Choi, 2018).

Yki has been shown to act through *thread*/*Diap1* to restrict tube size (Robbins et al., 2014) and absence of Yki results in reduced transcription of *Diap1.* While additional mechanistic evidence of the role of Diap1 in actin polymerization and growth control remains to be gathered, earlier studies have reported a function of Diap1 in cell migration, independent of its role in apoptosis. The authors showed that border cells of the *Drosophila* follicle epithelium lacking *Diap1* have low levels of F-actin and actin binding proteins and thus fail to migrate (Geisbrecht and Montell, 2004). It will be interesting to explore in future studies whether the defects in tracheal tube length of *yki* mutant embryos are due to an additional role of *Diap1* in the regulation of the actin cytoskeleton.

Based on our results, we propose a model in which Yki regulates development of the tracheal tubes by at least two mechanisms (Fig. 12). First, apical Yki binds to Tsr and facilitates the docking of a pool of Yki molecules to the apical actin cortex, thereby limiting its nuclear localization, a requirement for transcription of target genes, such as *diap1* (Fig. 12A). These findings together with previous observations in wing imaginal disc (Xu et al., 2018) point to a transcription-independent role of Yki in the apical cell cortex to regulate the apical cytoskeleton. Second, residual Yki molecules, which do not bind Tsr, are free to translocate to the nucleus and transcribe *Alas* to regulate crosslinking of cuticular proteins, and genes involved in in tissue size regulation, such as *diap1* (Fig. 12A). Thereby, *yki* not only regulates tissue size, but also contributes to the establishment of an extracellular barrier necessary for tissue tightness and tracheal gas filling.

**Figure 12.**
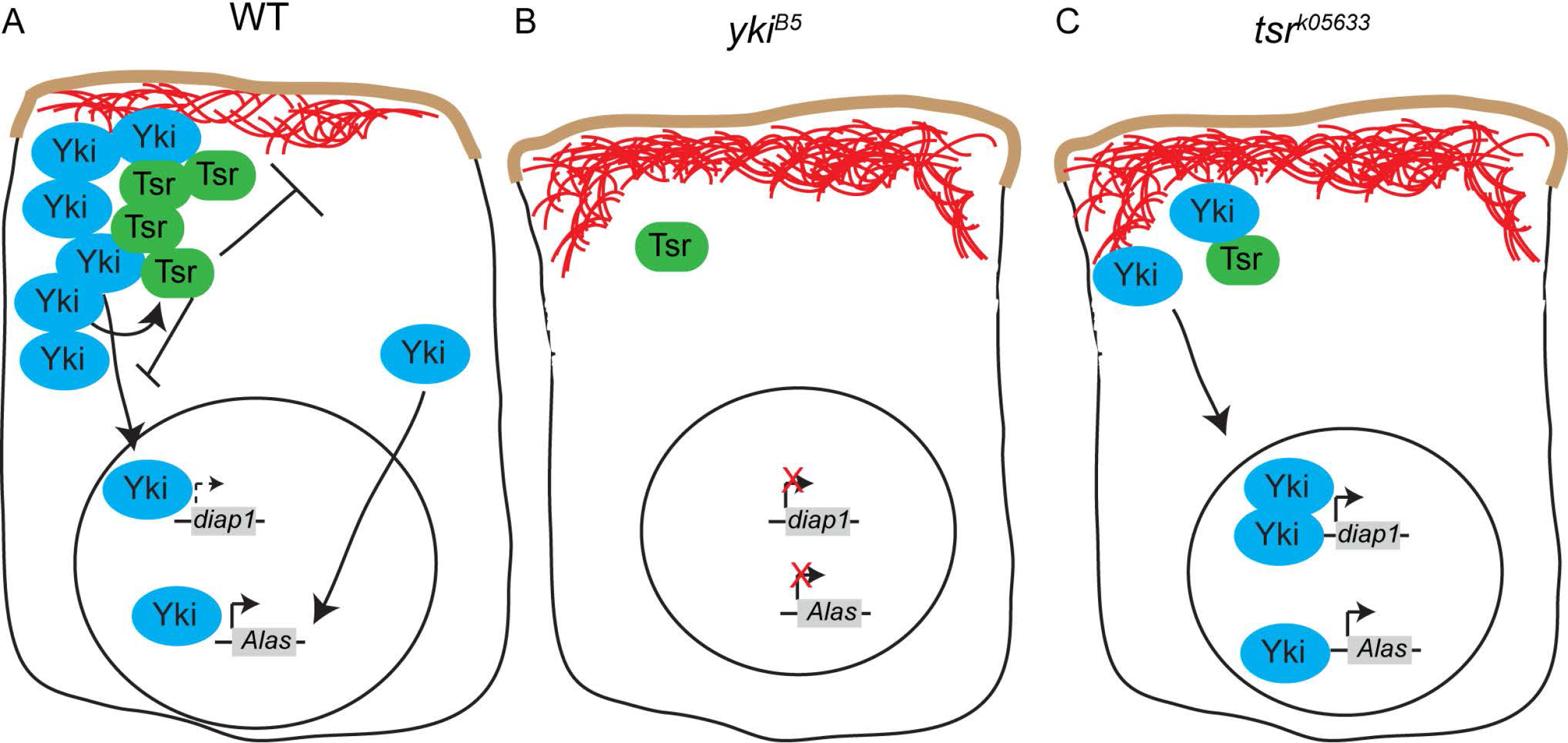
Model to explain how Yki and Tsr together control tracheal tube growth. (**A**) Yki and Tsr cooperate in the apical cell cortex to regulate membrane size and subsequently tissue growth. Tsr is a negative regulator of Yki nuclear translocation. Only a small portion of Yki is able to localize to the nucleus and to transcribe Yki-target genes necessary for tissue growth (e.g. *diap1*). Yki also transcribes genes required for tissue water-tightness and gas filling (e.g. *Alas*). Actin is marked in red. Arrow indicates high transcription. Dashed arrow indicates low transcription. (**B**) When Yki is absent, Tsr protein levels are reduced, resulting in increased apical F-actin and membrane growth. Yki target genes for tissue growth and water-tightness are no longer transcribed, resulting in longer tubes with defects in gas-filing. Actin is marked in red. Red cross indicates absence of transcription. (**C**) In the absence of Tsr, Yki protein levels are reduced and not maintained apically, allowing Yki molecules to translocate to the nucleus, resulting in stronger *diap1* transcription. However, higher Diap1 levels do not account for abnormal tube elongation. Rather, F-actin accumulates apically and apical membrane growth is increased, leading in longer tubes. Actin is marked in red.

Absence of Yki lowers Tsr levels and, therefore results in increased apical F-actin accumulation (Fig. 12B). Furthermore, absence of Yki prevents transcription of *Diap1* and *Alas*, thus giving rise to longer and water-permeable tubes. The tracheal growth phenotype of *yki* mutants can be partly be attributed to increased apical F-actin (via reduction of apical Tsr) and partly to the absence of *Diap1* transcription, since expression of *Diap1* in *yki* mutants only partially rescues the tube elongation phenotype.

Similarly, in the absence of *tsr*, the total levels of Yki are decreased but higher amount of Yki molecules translocate to the nucleus to transcribe higher levels of Yki-target genes, e.g. *Alas* and *Diap1* (Fig. 12C). At the same time, apical F-actin accumulates. We, therefore, propose that increased apical F-actin, rather than increased expression of *Diap1* leads to over-elongation of the tracheal tubes in *tsr* mutants. Tsr must also be signaling through F-actin to control tube length independently of the Yki/Diap1 pathway. Three lines of evidence favor this hypothesis: i) *yki, tsr* double mutants show more severe tube length defects as compared to single mutants, indicating that Tsr does not only signal through Yki/Diap1 (Fig. 6 D, E), ii) tracheal expression of *yki* in *tsr* mutants only partially restores the tube length, and iii) tracheal expression of Diap1 partially rescues *yki* mutants (Fig. 1, E), but not *tsr* mutants (data not shown).

Similarly, absence of Yki lowers Tsr levels and, therefore results in increased apical F-actin accumulation (Fig. 12C). Furthermore, absence of Yki prevents transcription of *Diap1* and *Alas*, thus giving rise to longer and water-permeable tubes. Thus, the tracheal growth phenotype of *yki* mutants can be partly be attributed to increased apical F-actin (via reduction of apical Tsr) and partly to the absence of *Diap1* transcription.

Taken together, our data uncover a dual role of Yki in tracheal development. Yki is required for proper gas-filling and tube growth, two processes that seem to be uncoupled. Overexpression of *Alas* rescues only the gas filling defects, but not the tube convolution phenotype of *yki* mutants. Conversely, tracheal length is not altered in *Alas* mutants. These results provide the first mechanistic view on the role of Yki in tube length control, which is independent of its role in cell proliferation and apoptosis. Our results contribute to further our understanding of the link between cortical actin organization and apical Yki activity in growth regulation, a link that could also be of importance in the emergence and progression of human diseases.

## Materials and Methods

### Fly stocks

The following *Drosophila* stocks were used: *w; yki^B5^,FRT42D/CyO* (kindly provided by DJ Pan), UAS-yki.GFP 4-9-Y (3^rd^ chromosome, kindly provided by Kenneth Irvine), *w; Mtf^ex234^/TM6C,dfd-YFP* (Tiklova et al., 2010), yw; *P{w[+mC]=lacW}Diap1[j5C8]/TM3,Sb* (BDSC #12093), *yw; P{w[+mC]=lacW}tsr[k05633]/CyO* (BDSC #12201), *yw; P{w[+mW.hs]=FRT(w[hs])}G13 tsr[N96A]/CyO* (BDSC #9108), *hpo^42-48^/CyO* (kindly provided by D.J. Pan), *w*; sp*/*Cyo*; Utr::GFP, Sqh::mCherry/*TM3* (kindly provided by Adam Martin), w*; *snaSco/CyO; P{UASt-Lifeact-RFP}* (BDSC #58362). For rescue experiments, *w; P{w[+mC]=UAS-DIAP1.H}* (BDSC #6657) *w; P{yw[+mC]=UAS-yki.V5.O}*attP2 (BDSC #28819), *yw; P{w[+mC]=UAS-tsr.N}/TM6B,Tb* (BDSC #9235) were used. Crosses for ectopic expression using *btl*-GAL4, w; *en*-Gal4,UAS.GFP (kindly provided by Georg Halder) were performed at 29°C. In all experiments CyO, TM3, and TM6C balancer strains carrying YFP transgenes were used to identify embryos of the appropriate genotypes.

### Generation of transgenic flies

To generate the UAS-Alas transgenic line, the *alas* cDNA (FI09607; obtained from DGRC) was cloned into pJFRC-MUH vector and injected into VK33 fly strain. UAS-Vc.yki (Vc is the C-terminal fragment of GFP) and UAS-Vn.Tsr (Vn is the N-terminal fragment of GFP) were generated by cloning of *yki* and *tsr* cDNA (LD21311 and LD06785 obtained from DGRC) into the pJFRC-MUH vector and injected into VK33 and attp40 fly strains, respectively.

### Cas9 genome engineering of *yki*

An N-terminal mKate2-Yki knock in chromosome was generated by CRISPR-Cas9 genomic editing using the “Scarless gene editing” design (flycrispr.molbio.wisc.edu), introducing and excisable TTAA-3xP3-dsRed-TTAA cassette originally removed by the PiggyBac transposase (Bruckner et al., 2017). The donor vector was modified to be resistant to Cas9 cleavage and bear the mKate2 and dsRed cassettes. Supplement 3 provides the sequences of the gRNA, its modified version with synonymous mutations in the donor vector and the complete donor vector repair template.

### Septate junction permeability assay

Stage 16.4 embryos (15.5 h AEL) were injected with 10kDa Rhodamine-Dextran and dye diffusion across epithelia of the main tracheal trunk was monitored by confocal microscopy 1 hour (60 min) to 4 hour (240min) after injection. Images were acquired with Zeiss LSM 510 Meta and Zeiss LSM 880 confocal microscopes and processed with Fiji.

### Quantitative Real-Time RT PCR

Total RNA was extracted from 150 embryos using Ambion RNAqueous kit. cDNA was synthesized using the reverse transcription Master Mix (Applied Biosystems). Real-time PCR was performed using primer sequences of: a) Alas *forward primer-*ACGGAACGTCTCCTACCTGA b) *Alas reverse primer-* TGCAGGTAGTGTCCGAATTG c) *Duox* forward primer-AGAAAGCAAAAATCGAGTGC d) *Duox* reverse primer-CGGTCTGACTATACATTTTCTCATAA e) *tsr forward primer-* TGTGCGAAATAACCGACCAA f) *tsr reverse primer-*ACACCAGAAGCCATTTTTCCT g) *yki forward primer-* GCGCCTTGCCGCCGG h) *yki reverse primer-*GCTGGCGATATTGGA i) Diap1 *forward primer-*: TCGTCAAATCTCAAC j) *Diap1 reverse primer-* TGAAGTCGAAACTTG. The experiments were performed for each genotype in triplicate and each experiment was repeated at least three times. Actin probe was used to normalize the total mRNA levels.

### Immunohistochemistry

The following antibodies were used at the indicated dilutions: mouse anti-Coracle (1:100, DSHB), fluorescein-conjugated ChtB (1:500, New England Biolabs), guinea pig anti-Vermiform (1:500; (Tsarouhas et al., 2007)), mouse anti-Crb (Cq4; 1:10, Developmental Studies Hybridoma Bank), mouse anti-Hnt (1:100; Developmental Studies Hybridoma Bank), guinea pig anti-Yurt (1:1000; gift from Ulrich Tepass), mouse anti-Mega (1:100; gift from Reinhard Schuh), guinea pig anti-Contactin (1:1500; gift from Manzoor Bhat), mouse anti-Fasciclin III (1:10; Developmental Studies Hybridoma Bank), rabbit anti-Varicose (1:500; (Bachmann et al., 2008)), guinea pig anti-Gasp (1:800; (Tsarouhas et al., 2007)), rabbit anti-Tsr (1:400; this study), rabbit anti-Echinoid (1:1000; gift from Laura Nilson), guinea pig anti-Uif (1:20; gift from Robert Ward), rabbit anti-Perlecan (1:1000; gift from Stefan Baumgartner), anti-Dityrosine (1:200, Japan Institute for the Control of Aging, Cat# MDT-020P), anti-Yki (1:200 for IF, 1:1000 for western blots; gift from Ken Irvine). Wheat germ agglutinin (WGA) (1:200; Thermo Fischer Scientific), chitin binding probe-633 (1:20; gift from Maria Leptin). Secondary antibodies conjugated to Cy3 (Jackson Immunochemicals) or Alexa Fluor 488, 568 and 633 (Molecular Probes) were used at 1:400 dilution. Images were obtained with Zeiss LSM 880 and Zeiss LSM 510Meta confocal setup and processed with Fiji.

### Cell number counting

Stained embryos were imaged with a laser-scanning confocal microscope (LSM780, Carl Zeiss) using a 63/1.4NA C-Apochromat oil-immersion objective. Individual Z stacks were taken with a step size of 0.2–0.5 μm and with a total Z sampling distance of 20-30 μm. The cell number counting was based on the nuclei numbers. To separate nuclei closely adjacent to each other (and potentially yielding false negatives), a 3D Watershed segmentation method utilized. Z-stacks binarization, Watershed digital image reconstructions and nuclei counting computed by the ImageJ/Fiji software.

### Proximity ligation assay (PLA)

PLA recognizes the potential interaction of endogenous proteins using antibodies to detect proteins in close proximity to one another (<40 nm). Third instar larva wing discs were dissected and fixed as described above. Primary antibodies against Tsr (rabbit anti-Tsr) and V5 epitope (mouse anti-V5, Invitrogen) were added and incubated overnight at 4°C. The Duolink PLA Kit (Sigma Aldrich) was used to incubate the tissue with the PLA probes PLUS and MINUS at 37°C for 1 hour. Ligation of the PLA oligonucleotides and amplification were performed at 37°C for 30 min and 100min, respectively. Samples were mounted in Duolink mounting media and imaged using Zeiss LSM 880.

### Antibody generation

Full length *tsr* of GenBank nucleotide sequence RE04257 was cloned into pGEX4T2 (N-terminus GST) vector using the following primers: *forward primer*-(BamHI) <u>CTCGGATCC</u>GCTTCTGGTGTAACTGTGTCTG and *reverse primer*-(EcoRI) <u>CTCGAATT</u>CTTATTGGCGGTCGGTG. The protein was expressed in *E.coli* BL21 cells, solubilized in 50mM Tris-HCl pH=8 and purified using GST Sepharose beads. Purified protein was dialyzed against PBS and used for antibody generation in rabbits at the Charles River Laboratories International, Inc. The rabbit anti-Tsr serum was purified by affinity purification and stored in 50% glycerol.

### Immunoprecipitation

Fly embryos of the following genotypes: *w^−^; btl-Gal4/Cyo,DfdYFP;UAS-yki.V5* and *w^−^; btl-Gal4/Cyo,DfdYFP* (as control) were used. Embryos were homogenized on ice using Dounce tissue grinder in 1mL of lysis buffer containing 130mM NaCl, 50mM Tris-HCL pH=8, 0,5% Triton-X and protease inhibitor (Roche). After 30min at 4°C under rotation the homogenate was centrifuged for 20min at 14.000rpm. Supernatant was incubated with the antibody for 2 hours. In the meantime 50μl of Protein G were washed 3 times with blocking solution and incubated with the antibody solution overnight at 4°C. Protein G beads were collected by centrifugation for 2 min at 3.000rpm and washed 4 times with lysis buffer. Beads were resuspended in 1.5X SDS sample buffer and heated for 5min at 95°C.

### Western Blot

Samples were separated by SDS-PAGE and blotted onto nitrocellulose 0.45 membrane (Amersham). After blocking in 5% BSA +TBST, the membrane was incubated overnight with rabbit anti-Tsr diluted 1:4000, anti-Yki diluted 1:1000 and anti-alpha-Tubulin diluted 1:5000 in blocking buffer. Peroxidase antibodies were used for detection.

### Cell Culture and Transfection

*Drosophila* S2R^+^ cells were cultured at 25 °C in Schneider’s *Drosophila* medium (Sigma) supplemented with 10% fetal bovine serum. Transfection of p*Act5*-Gal4 and Vn-Yki into S2R^+^ cells was performed with FuGENE HD (Promega) according to the manufacturer’s protocol. After 48 h, cells were treated with 100μg/ml cycloheximide for the indicated times. Cells were harvested, washed with ice-cold PBS (120 mM NaCl in phosphate buffer at pH 6.7), resuspended in lysis buffer (containing 10% glycerol; 1% Triton X-100; 1.5 mM MgCl_2_; 120 mM NaCl; 100 mM PIPES, pH 6.8; 3 mM CaCl_2_; 1 mM PMSF and Complete^TM^). Cells were lysed on ice for 20min and lysates were centrifuged at 14.000 rpm for 20min at 4°C. Sample buffer 3x SDS was added to supernatant and boiled for 5min at 95°C.

### Mass spectrometry analysis

Lysates of stage 17 embryo expressing a tracheae-specific Yki-V5 construct were immunoprecipitated (described above) using the mouse anti-V5 antibody (Invitrogen). After electrophoretic separation on SDS gel and Coomassie staining, lanes were cut into 10 slices each, digested in-gel with trypsin, and subjected to GeLC MS/MS analysis.

GeLC MS/MS analysis was performed on an Ultimate3000 nanoLC system interfaced on-line to a LTQ Orbitrap Velos hybrid tandem mass spectrometer (both Thermo Fisher Scientific, Bremen, Germany). Internal standard (GluFib peptide) was spiked into each sample prior analysis. Proteins were identified by Mascot software v.2.2.04 (Matrix Sciences Ltd, London, UK) by searching against Drosophila protein sequences in NCBI database (May 2014, 231613 entries) under the following settings: 5ppm and 0.5Da mass accuracy for precursor and fragment ions, respectively; enzyme specificity – trypsin; maximal number of allowed miscleavages – two; variable modifications – methionine oxidation, N-terminal protein acetylation, cysteine propionamide. The result of the database search was evaluated by Scaffold software v. 4.3.2 (Proteome Software, Portland) using 95% and 99% probability threshold for peptides and proteins respectively; minimal number of matched peptides was set on two. Calculated False Discovery Rate (FDR) (standards Scaffold feature) for peptides and proteins was below 0.5%. Relative quantification of proteins was performed using MaxQuant software; absolute values of individual proteins were normalized on total protein intensity and internal standard.

### Electron microscopy

Embryos and larvae were fixed in 2% Glutaraldehyde in 0.1M PB buffer pH=7.2 for 20min at room temperature. Embryos were hand devitellinized. Both embryos and larvae were transferred in microcentrifuge tubes and fixed in 1%OsO_4_/2% Glutaraldehyde and then 2% OsO_4_. Specimens were washed and dehydrated in Araldite. Ultrathin sections of 0.1μm were prepared and analyzed with Tecnai 12 BioTWIN (FEI Company).

### Fluorescence Microscopy Imaging of live imaginal discs and FCS

Fluorescence imaging and FCS measurements were performed on two uniquely modified confocal laser scanning microscopy systems, both comprised of the ConfoCor3 system (Carl Zeiss, Jena, Germany) and consisting of either an inverted microscope for transmitted light and epifluorescence (Axiovert 200 M); a VIS-laser module comprising the Ar/ArKr (458, 477, 488 and 514 nm), HeNe 543 nm and HeNe 633 nm lasers and the scanning module LSM510 META; or a Zeiss LSM780 inverted setup, comprising Diode 405 nm, Ar multiline 458, 488 and 514 nm, DPSS 561 nm and HeNe 633 nm lasers. Both instruments were modified to enable detection using silicon Avalanche Photo Detectors (SPCM-AQR-1X; PerkinElmer, USA) for imaging and FCS. Images were recorded at a 512×512 pixel resolution. C-Apochromat 40x/1.2 W UV-VIS-IR objectives were used throughout. Fluorescence intensity fluctuations were recorded in arrays of 10 consecutive measurements, each measurement lasting 10 s. Averaged curves were analyzed using the software for online data analysis or exported and fitted offline using the OriginPro 8 data analysis software (OriginLab Corporation, Northampton, MA). In either case, the nonlinear least square fitting of the autocorrelation curve was performed using the Levenberg-Marquardt algorithm. Quality of the fitting was evaluated by visual inspection and by residuals analysis. Control FCS measurements to asses the detection volume were routinely performed prior to data acquisition, using dilute solutions of known concentration of Alexa488 dye. The variability between independent measurements reflects variability between cells, rather than imprecision of FCS measurements. For more details on Fluorescence Microscopy Imaging and FCS, refer to Supplement 2.

## Acknowledgments

We are grateful to Stefan Luschnig, Ulrich Tepass, Reinhard Schuh, Alan Fanning, Manzoor Bhat, Bernard Mechler, Ken Irvine, Stefan Baumgartner, Robert Ward, Maria Leptin, Laura Nilson and Developmental Studies Hybridoma Bank (DSHB; Iowa) for sharing antibodies. Adam Martin, Duojia Pan, Ken Irvine, Georg Halder and the Bloomington *Drosophila* Stock Center for *Drosophila* strains. The *Drosophila* Genomics Resource Center (DGRC; Indiana) for clones. We are indebted to the Mass Spectrometry Facility and especially to Anna Shevchenko, to the Antibody Facility (Patrick Keller), the Gene Expression Facility (Julia Jarrells), the Light and Electron Microscopy Facility (Jan Peychl and Weihua Leng) at MPI-CBG for the outstanding technical assistance. We also would like to thank Catrin Hälsig, Amelia Aragones-Hernandez and Michaela Burkon for their technical assistance and the Knust lab members for their fruitful discussions. K.S. was supported by a Wenner-Gren stipendium från Wenner-Gren Stiftelserna. The work at MPI-CBG was supported by the Max-Planck Society.

## Supporting information

**Suppl. Fig 1.**
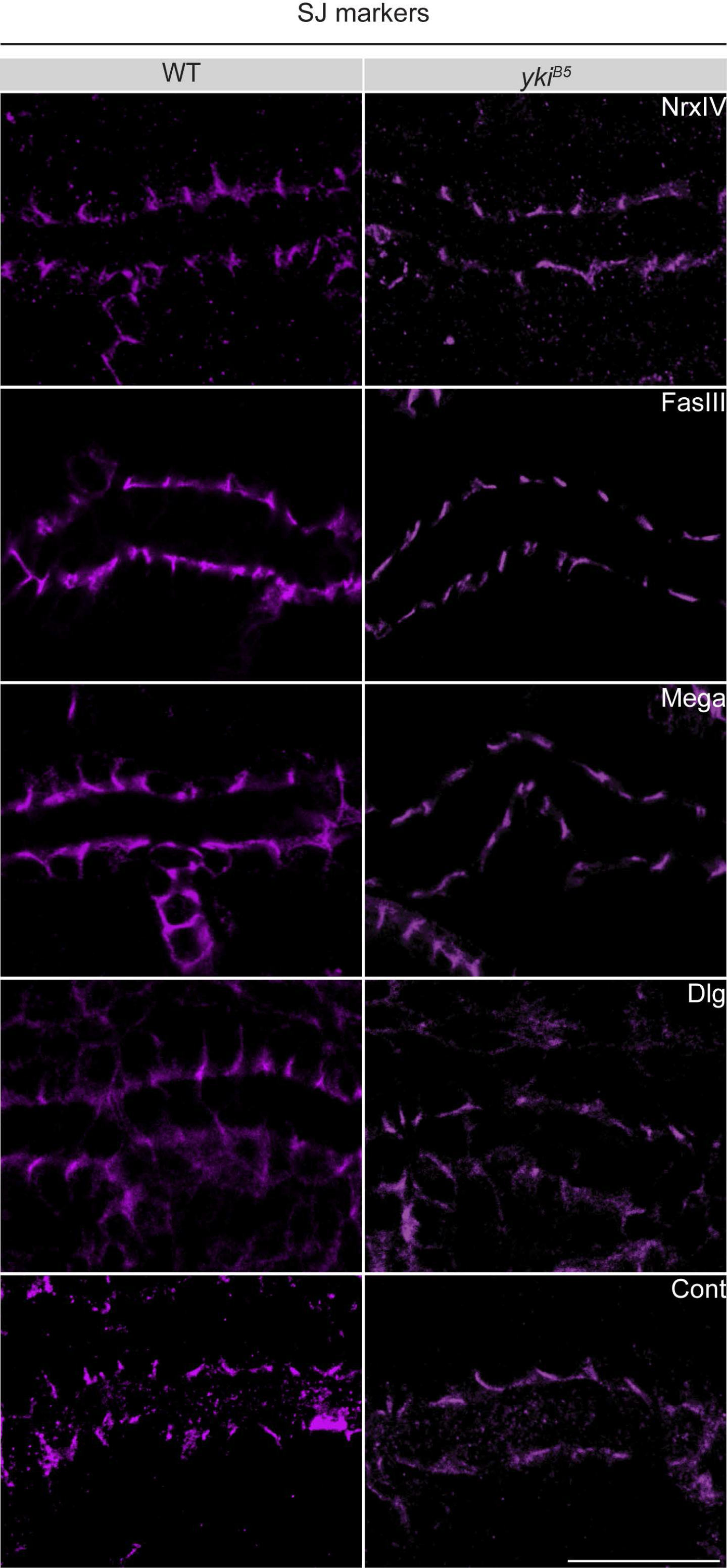
Localization of septate junction proteins is not compromised in the absence of Yorkie. Single confocal sections of early stage 17 dorsal trunks stained for septate junction (SJ) proteins NrxIV, FasIII, Mega, Dlg and Cont. In *yki*^*B5*^ mutant embryos SJ proteins do not change their expression or localization. Scale bar: 10 μm.

**Suppl. Fig 2.**
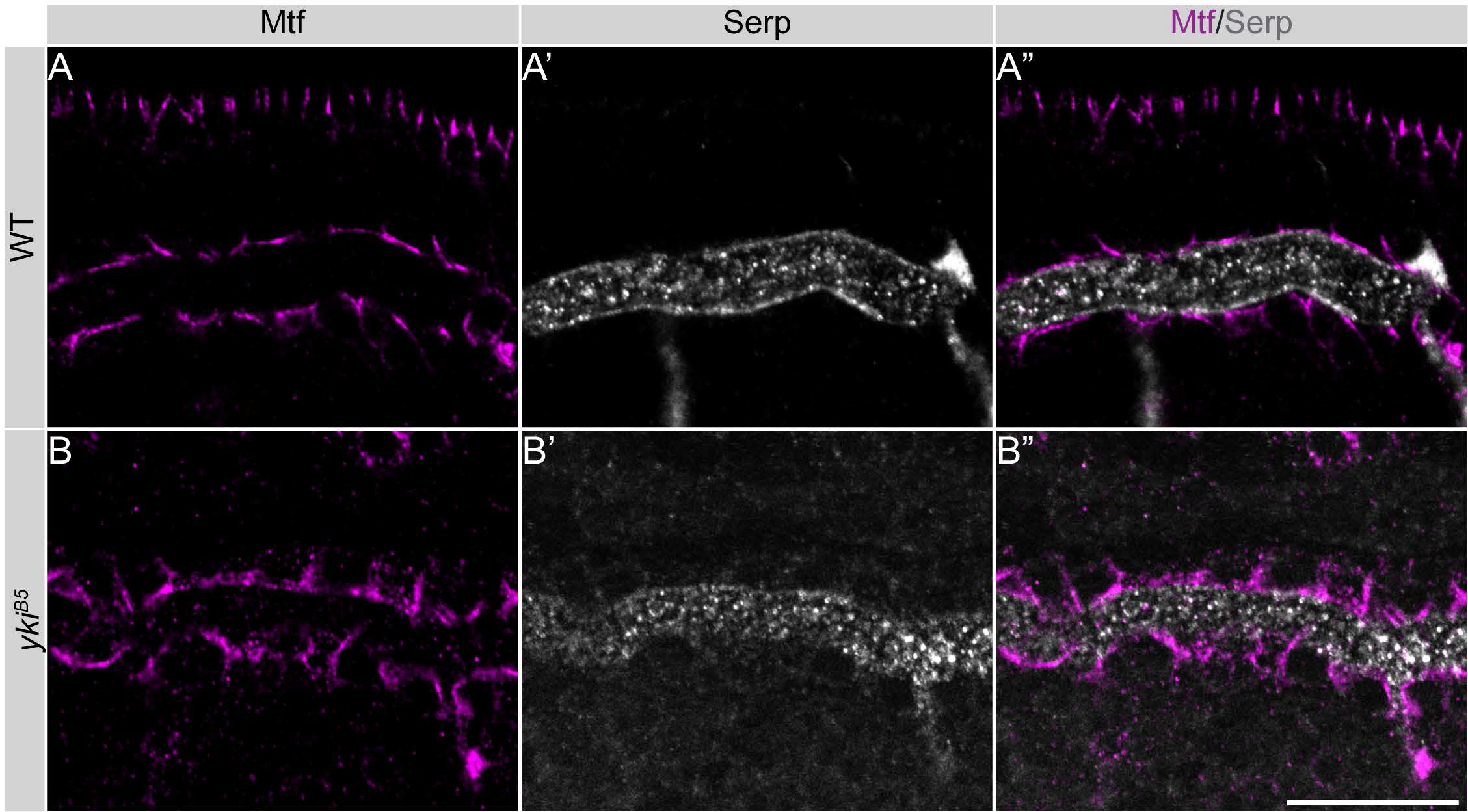
Localization of the luminal matrix protein Serp is not compromised in the absence of Yorkie. (A-B”) Single confocal sections of early stage 17 dorsal trunks stained for the septate junction protein Mtf (A, B and A”. B”) and the luminal matrix marker, Serp (A’, B’ and A”. B”). In *yki*^*B5*^ mutant embryos Serp does not change its expression or its localization. Scale bar: 20 μm.

**Suppl. Fig 3.**
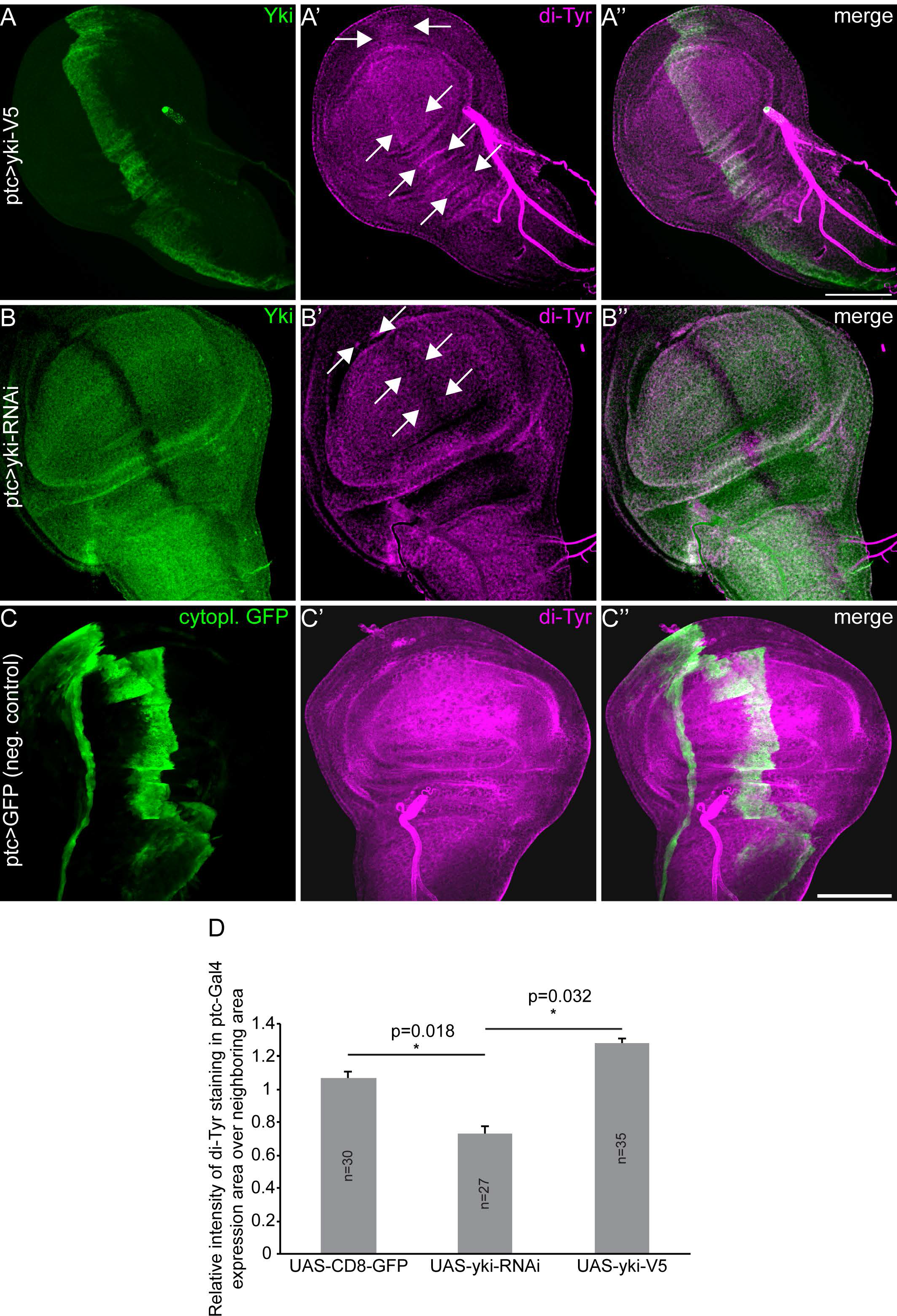
Yorkie levels influence Dityrosine network in wing discs. (**A-C”**) Yki overexpression (A-A”) or knockdown (B-B”) by *ptc*-Gal4 leads to increased or decreased levels of dityrosine staining in the A/P boundary of third instar wing discs, respectively. GFP (ptc>GFP) was used as a negative control (C-C”). Scale bar: 100μm. (**D**) Quantification of relative intensities of dityrosine in the *ptc*-Gal4 domain.

**Suppl. Fig 4.**
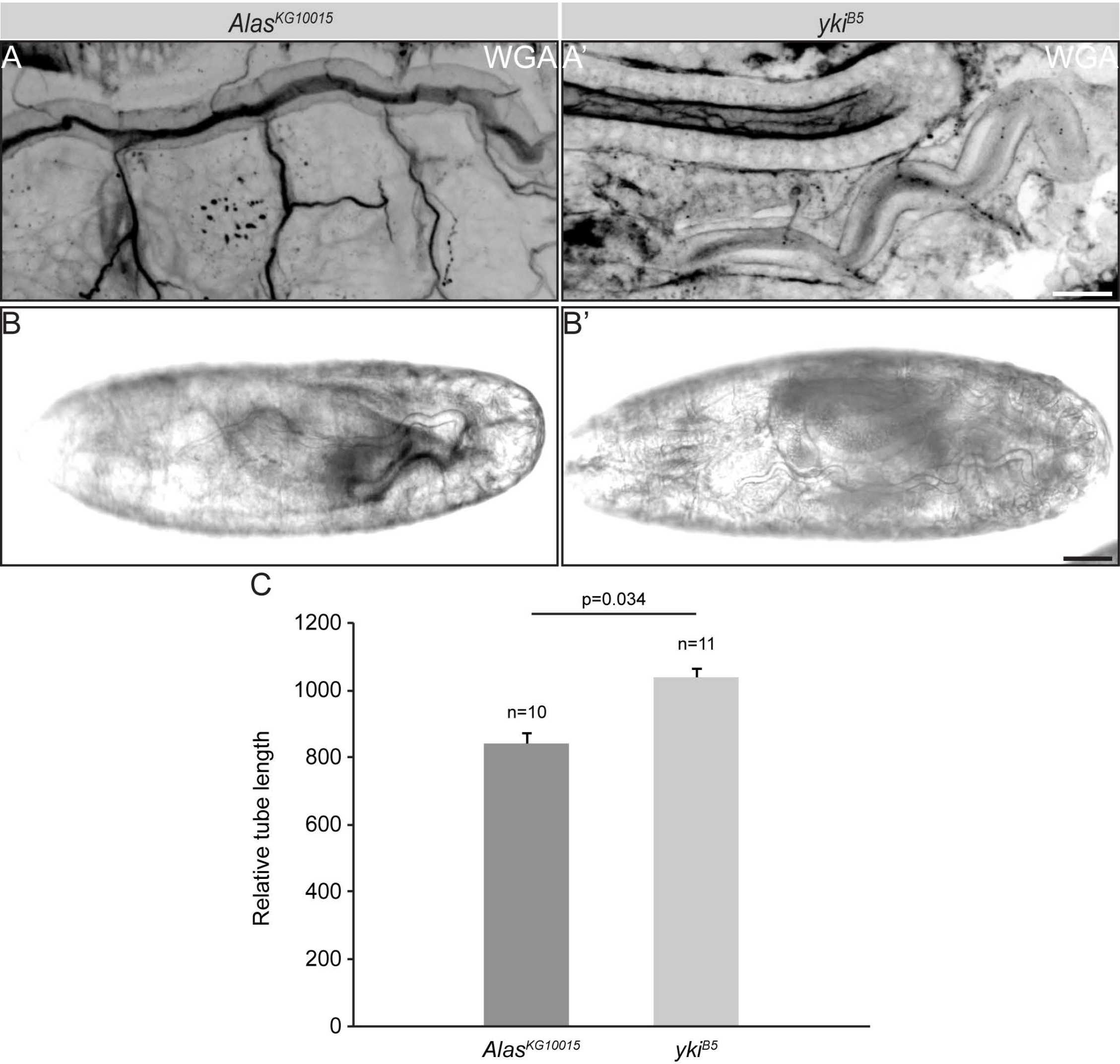
Loss of *Alas* does not affect axial tube elongation. (**A-B’**) Lateral views of *Alas*^*KG10015*^ (A) and *yki*^*B5*^ (A’) embryos stained with WGA, and of live embryos (B, B’) showing that tubes in *alas*^*KG10015*^ mutants are shorter than in embryos mutants for *yki* Scale bars: 20 μm (A, A’, B, B’). (**C**) Bar graph showing the relative tube length of *Alas*^*KG10015*^ and *yki*^*B5*^ embryos.

**Suppl. Fig 5.**
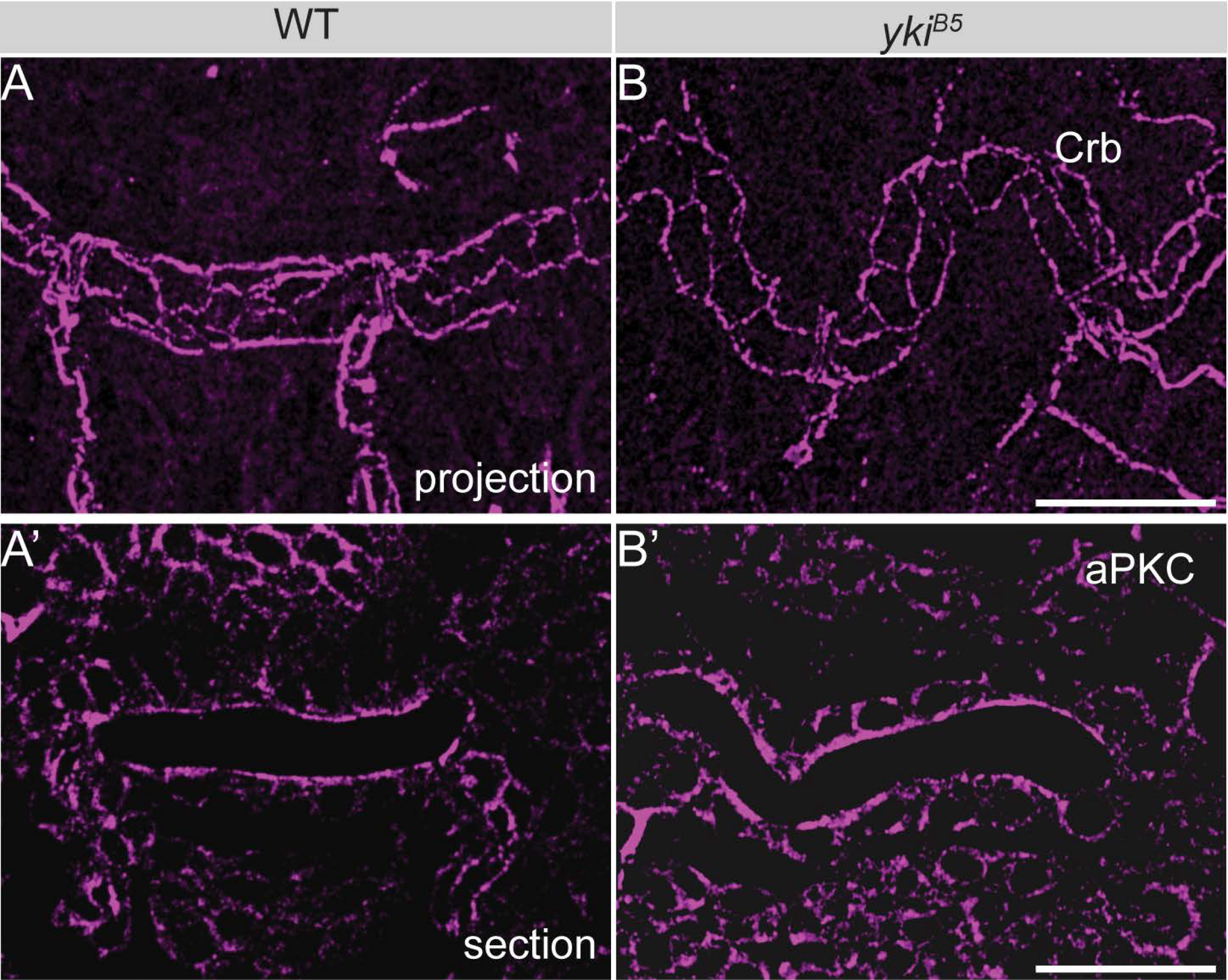
Cell polarity is preserved in the absence of Yorkie. (**A-B**) Confocal projections of the dorsal trunk metamere 8 of early stage 17 wild type (WT) and *yki*^*B5*^ embryos labeled for Crb. (**A’-B’**) Superficial sections of the dorsal trunk metamere 8 of early stage 17 wild type and *yki*^*B5*^ embryos labeled for aPKC. Scale bars: 20 μm.

**Suppl. Fig 6.**
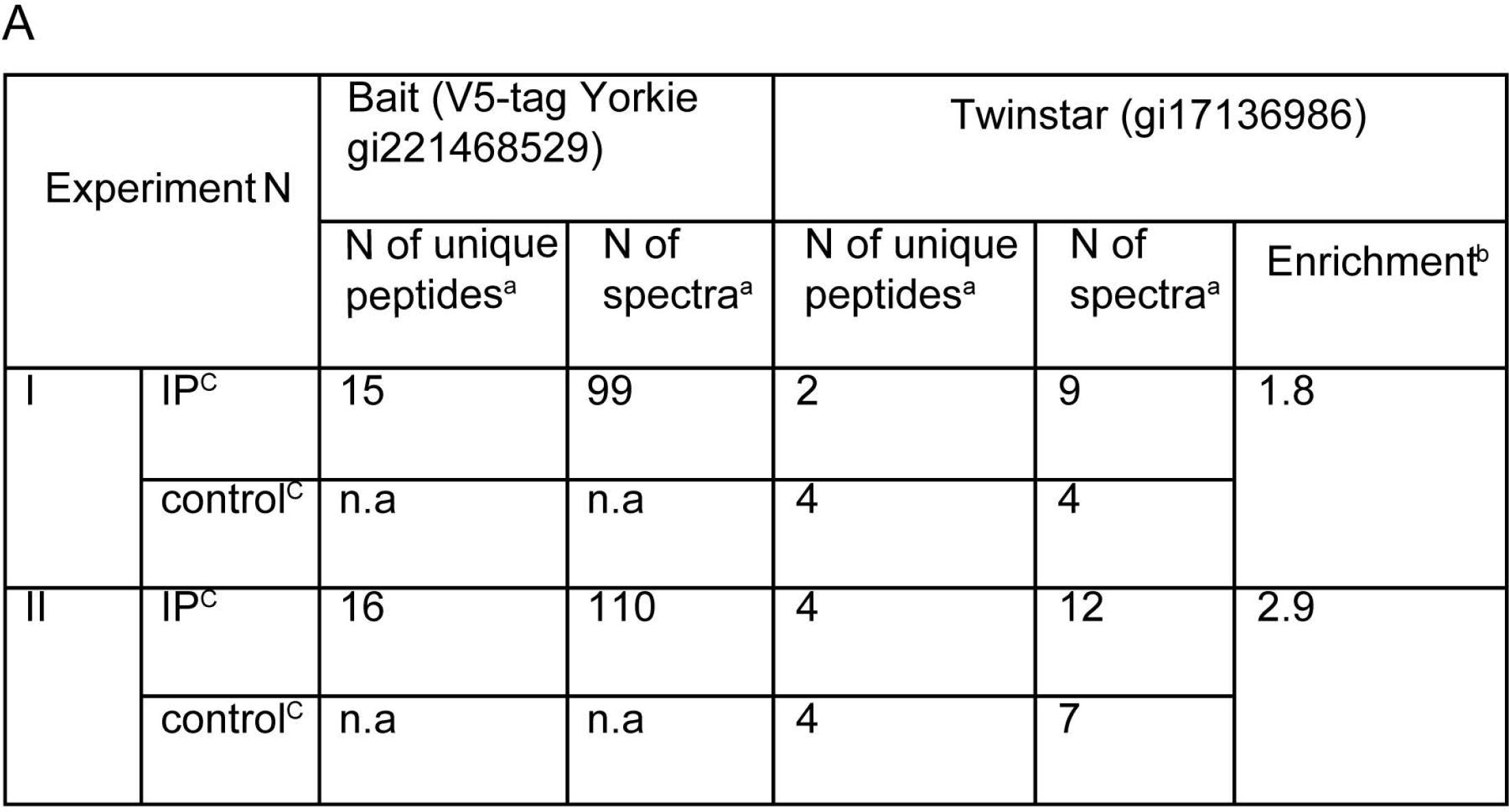
Twinstar and actin specifically associate with Yorkie. (**A**) Mass spectrometric identification and relative quantification of Twinstar protein in repetitive immunoprecipitations of V5-tagged Yorkie. a – as reported by Scaffold software under setting described in materials and methods section; b – calculated as ratio between relative normalized intensities in immunoprecipitation and corresponding control; Twinstar was not detected in the control of the experiment N1. c - immunoprecipitation of V5-tagged Yorkie and control experiments were performed as described in materials and methods section.

**Suppl. Fig 7.**
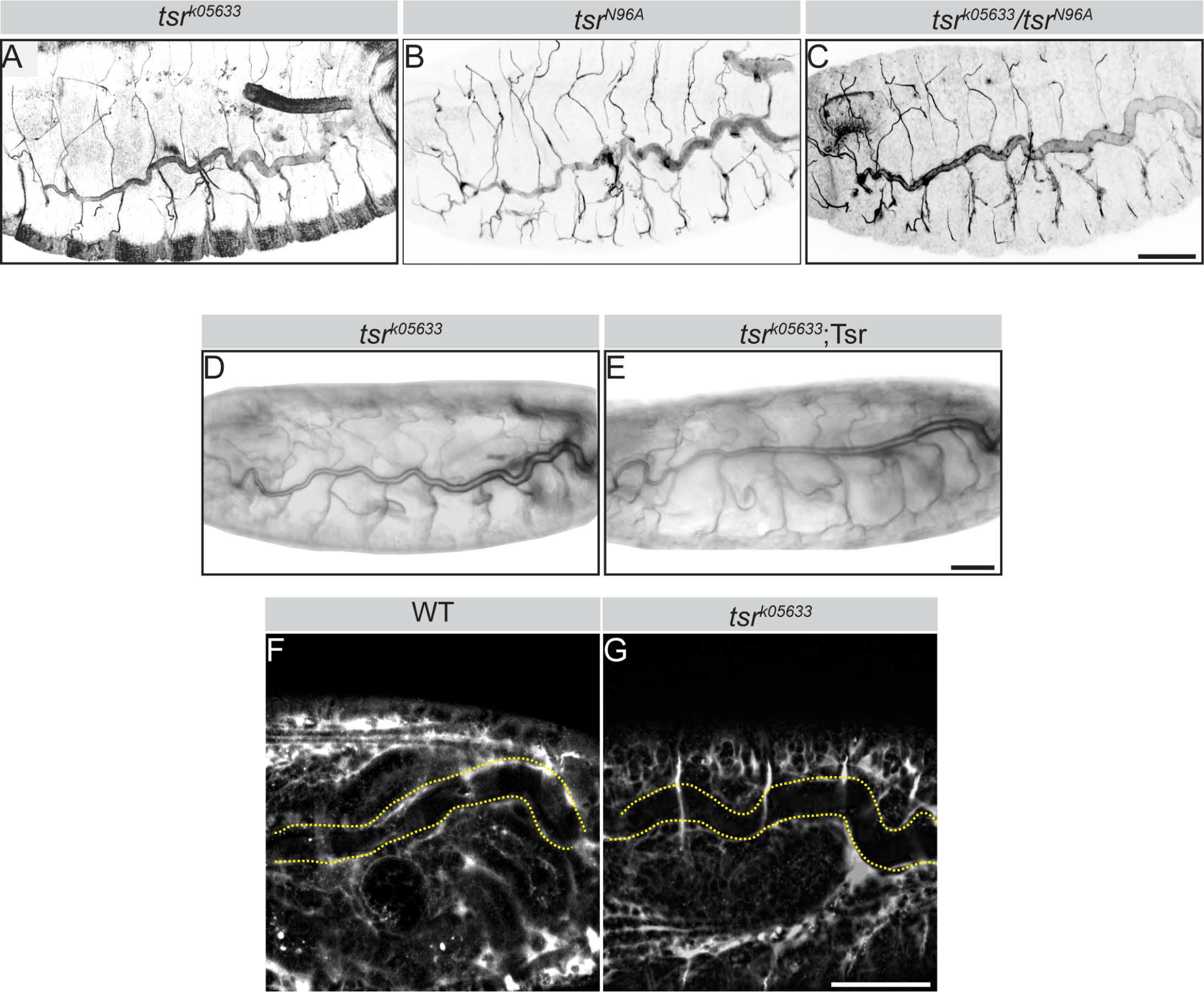
Twinstar mutants exhibit defects in tube size, but not gas-filling and paracellular barrier defects. (**A-C**) Tracheal phenotypes of two different alleles of *twinstar*, *tsr*^*k05633*^ and *tsr*^*N96A*^, stained for Verm. (**D-E**) Overviews of *tsr*^*k05633*^ stage 17 embryos, showing proper tracheal gas filling in the absence (D) or presence (E) of a transgene expressing a wild type Twinstar. Scale bars: 50μm (A, B). (**F-G**) Fluorescent 10kDa Dextran injected into the body cavity of WT (F) and *tsr*^*k05633*^ (G) mutants does not leak into the tracheal lumen (dashed lines), indicating that the paracellular barrier in *tsr*^*k05633*^ mutants is intact. Scale bar: 50μm.

**Suppl. Fig 8.**
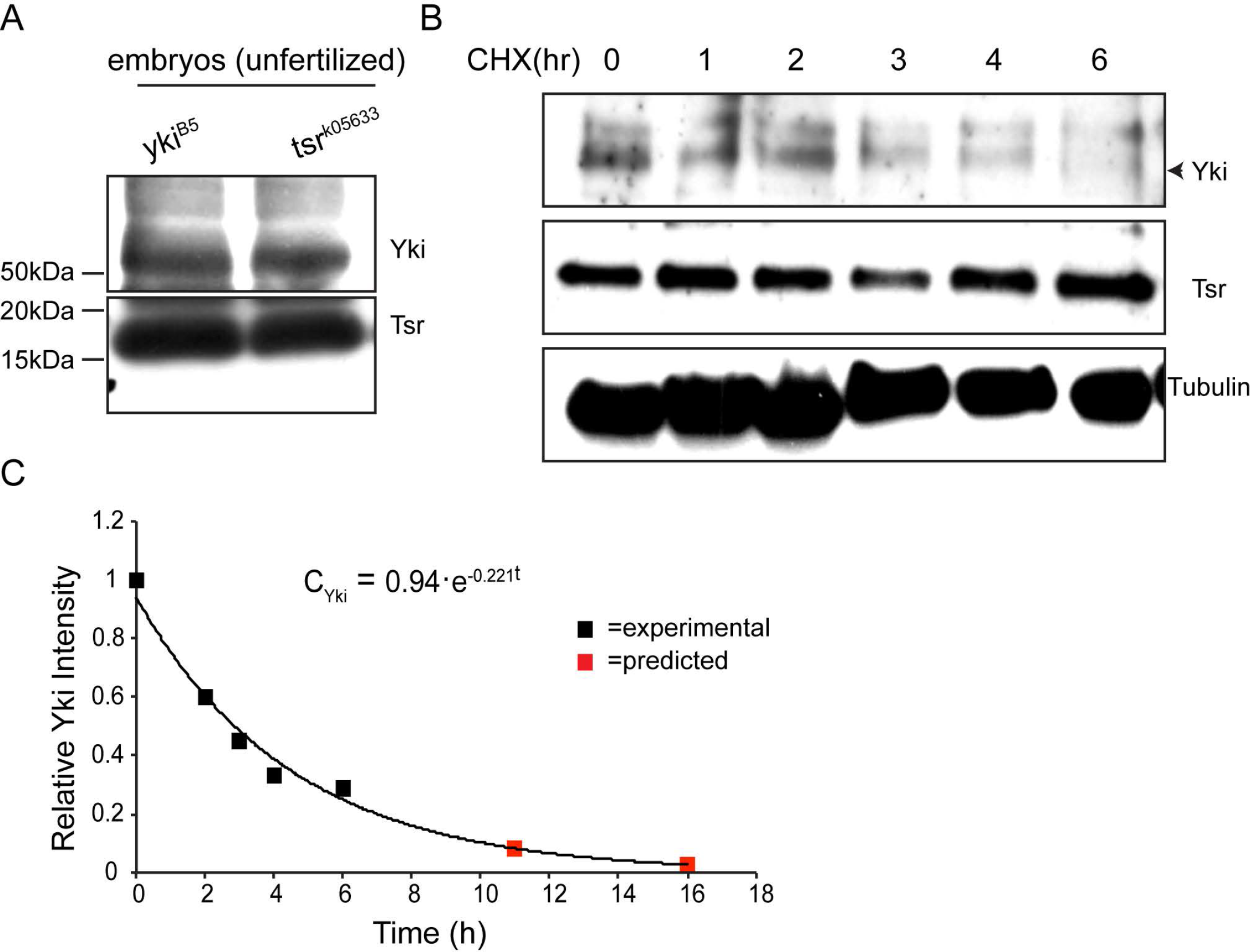
Yki and Tsr are maternally contributed. (**A**)Protein lysates from unfertilized eggs of *yki*^*B5*^ and *tsr*^*k05633*^ heterozygote females. (**B**)Western blots from cell lysates of S2 cells expressing Yki and Tsr, treated with cycloheximide (CHX) for 0, 1, 2….6 hours. Tubulin is used as a loading control. (**C**) Degradation kinetics (*N*(*t*) = *N*_0_ *e*^−*λt*^), derived from panel B, shows the level of Vn-Yki in S2 cells after protein synthesis inhibition by cycloheximide (CHX). 0, 1, 2…18 refer to hours after CHX addition. Data were collected from two independent experiments.

**Suppl. Fig 9.**
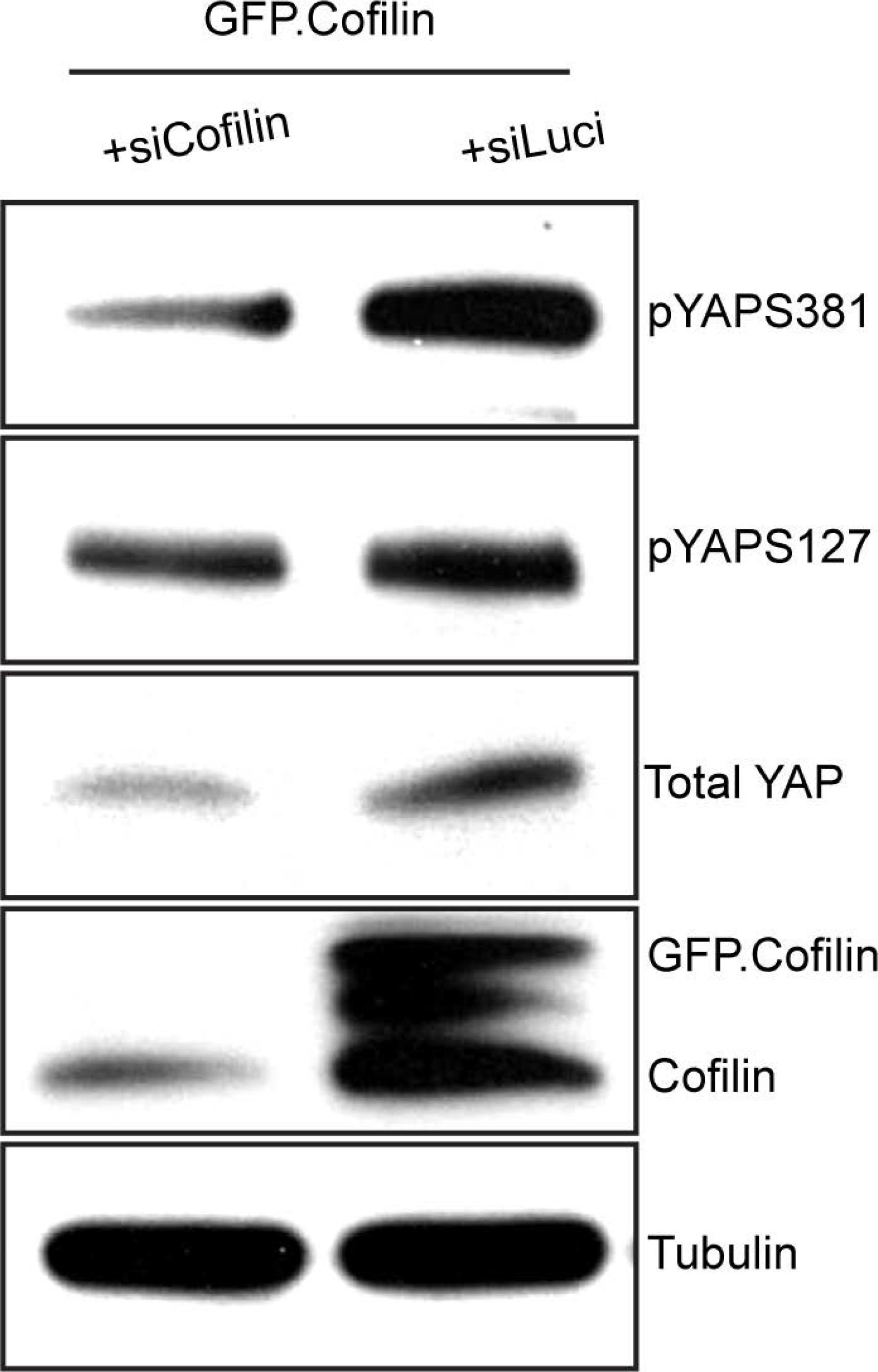
Knock-down of Cofilin reduces YAP protein levels and decreases its phosphorylation state. Cofilin knockdown in HEK293T cells expressing GFP-Cofilin reduces YAP total protein levels as well as YAP phosphorylation at S381 and S127. Tubulin is used as loading control.

**Suppl. Fig 10.**
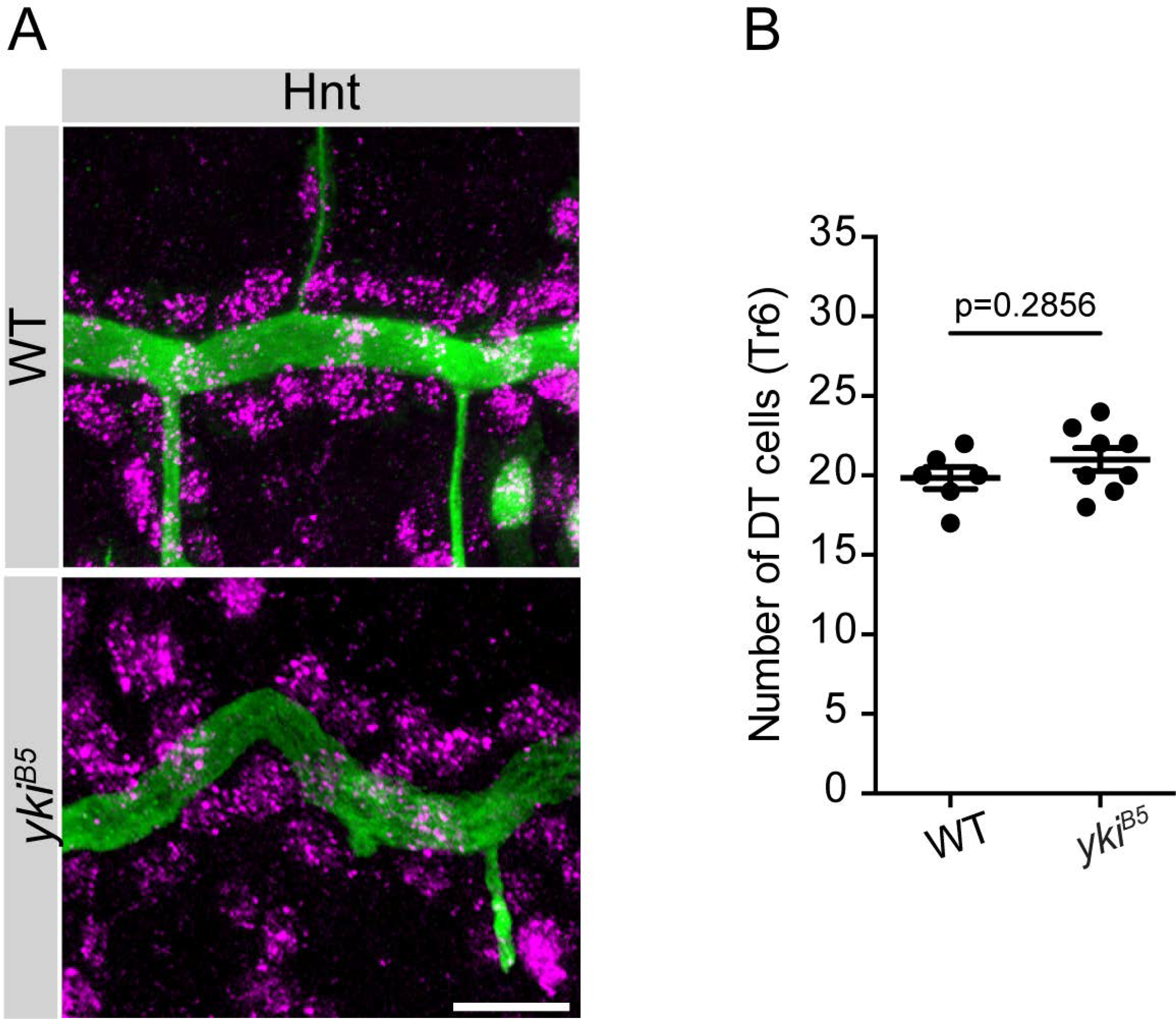
Tracheal cell number does not change in *yki*^*B5*^ mutant embryos. (**A**) Confocal projections showing the dorsal trunk (DT) of stage 17 wild type (WT) and *yki*^*B5*^ mutant embryos stained for the luminal protein Gasp (green) and the nuclear marker Hnt (Hindsight, magenta). Scale bars: 10μm. (**B**) Plot showing the average number of cells, which correspond to the number of marked nuclei, in wild type (WT) (*n* = 6) and *yki*^*B5*^ (*n* = 8) mutant embryos of the tracheal metamere 6 (Tr6) of stage 17 embryos. Values were calculated based on a 3D Watershed-based image transformation (see details in Materials and Methods). Unpaired two tailed *t*-test was performed to obtain the indicated *p* value. Error bars represents s.e.m.

**Suppl. Fig 11.**
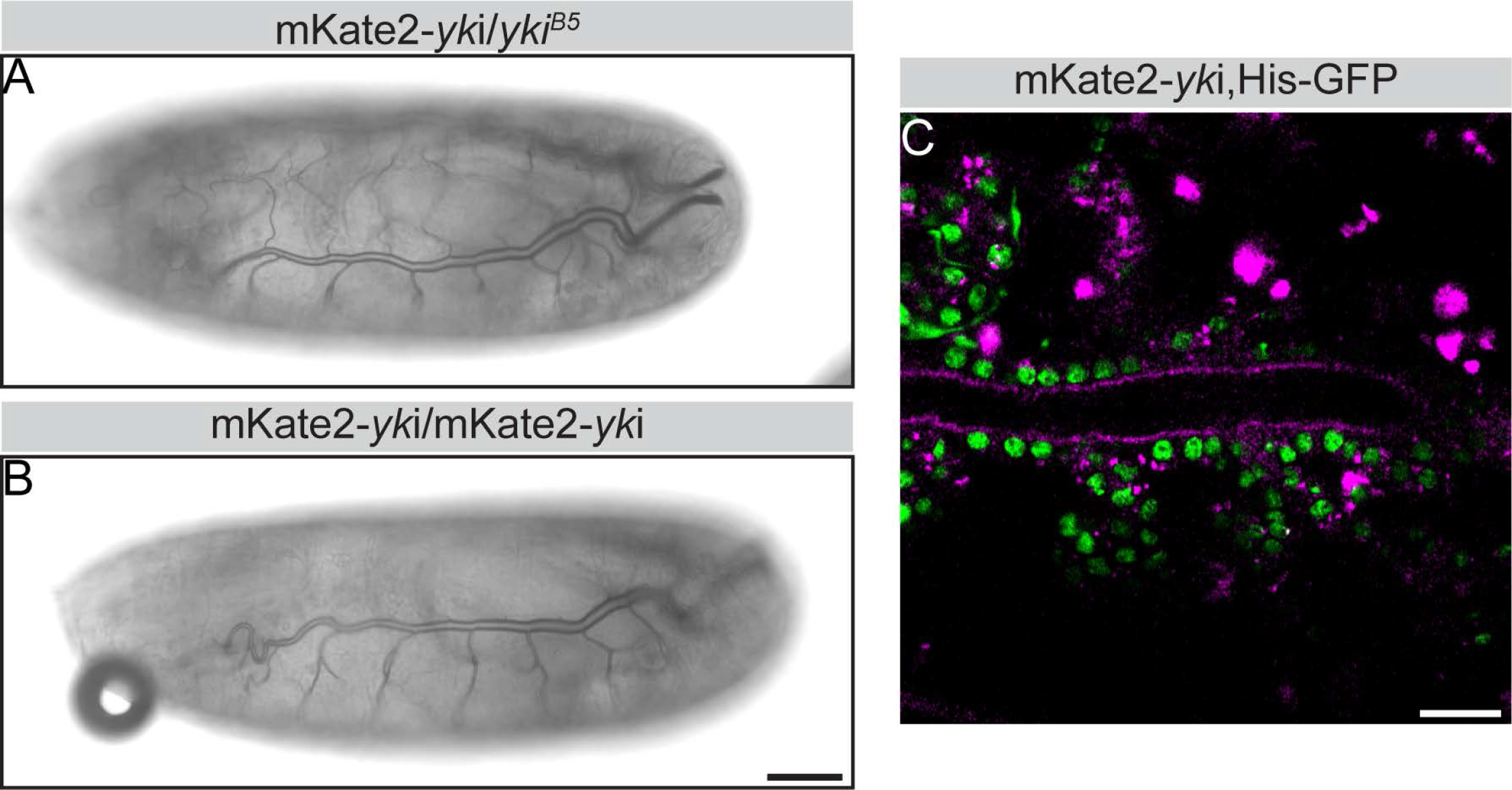
The endogenously tagged *yki* allele (mKate2-*yki*) is homozygous viable and rescues the mutation. (**A, B**) Lateral views of an mKate2-*yki* /*yki*^*B5*^ (A) transheterozygous and an mKate2-*yki* homozygous (B) stage 17 embryo showing normal tube length and gas-filling. Scale bar: 20 μm. (C) Confocal section of part of the DT of an mKate2-*yki* embryo of stage 17 expressing Histone-GFP (green). Yki (magenta) is localized apically in the tracheae, but cannot be detected in the nuclei (marked by GFP). Scale bar: 50 μm

### Supplement 1. Overview of actin peptides detected by mass spectrometry in immunoprecipitations of V5-tagged Yorkie and the corresponding controls

Peptides detected with mass spectrometry are highlighted with different colors: grey indicates peptides shared between all actin sequences (peptides in *italic* were used for relative quantification of total actin); yellow - peptides unique for Act5C/Act42A, blue – peptides unique for Act79B/Act88F/Act87E, green – peptides unique for Act57B.

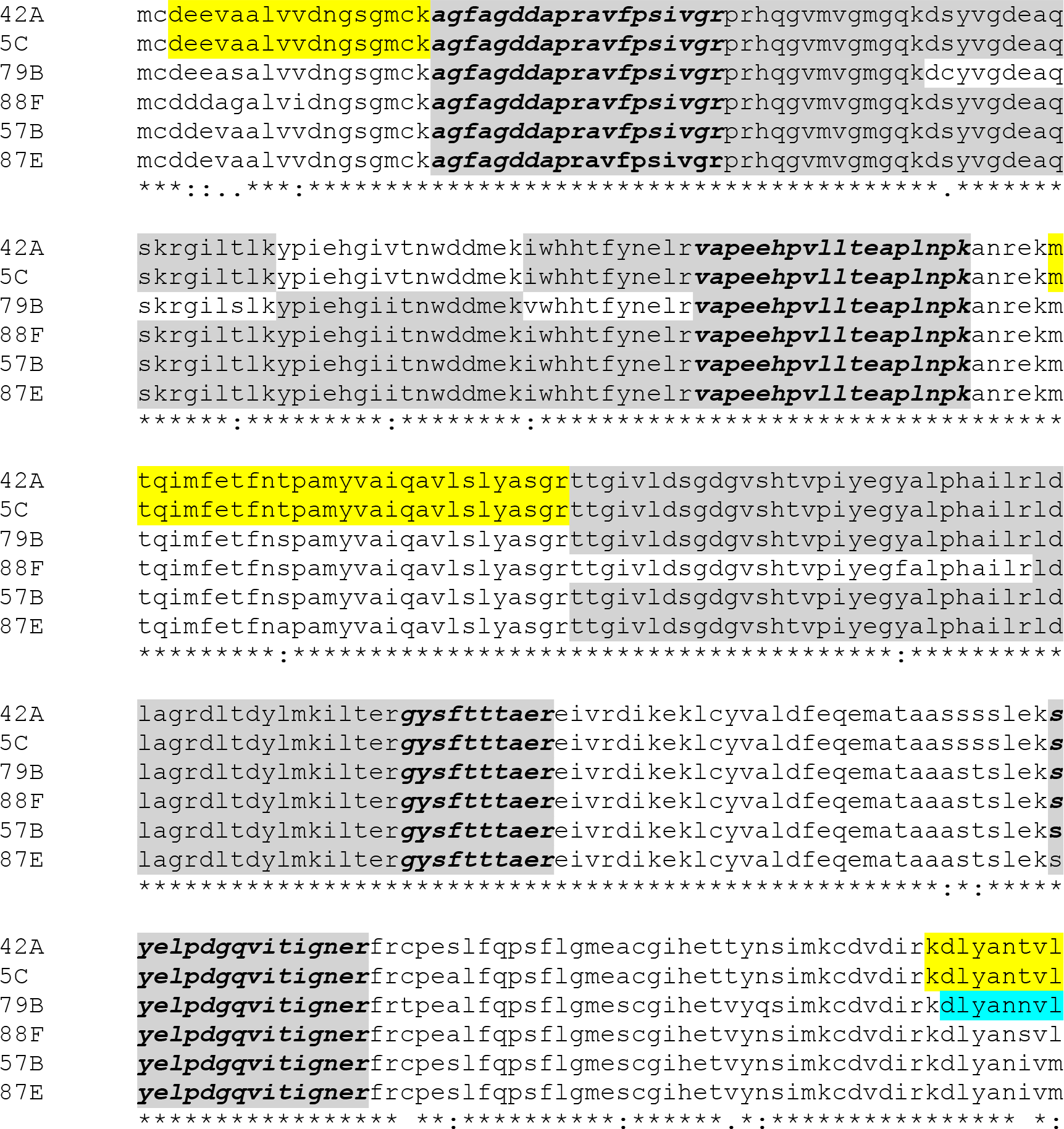

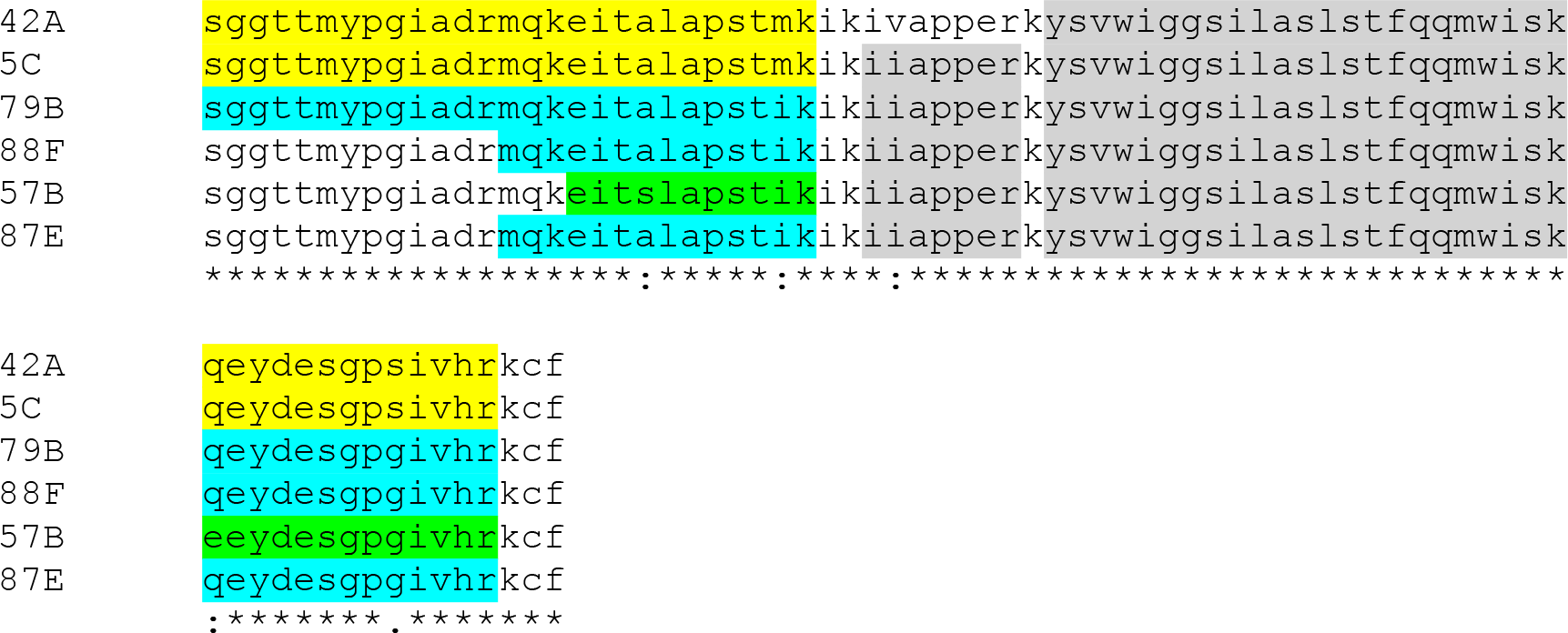

### Supplement 2. Background on Fluorescence Microscopy Imaging and FCS

Two individually modified instruments (Zeiss, LSM 510 and 780, ConfoCor 3) with fully integrated FCS/CLSM optical pathways were used for imaging. The detection efficiency of CLSM imaging was significantly improved by the introduction of APD detectors. As compared to PMTs, which are normally used as detectors in conventional CLSM, the APDs are characterized by higher quantum yield and collection efficiency – about 70 % in APDs as compared to 15 – 25 % in PMTs, higher gain, negligible dark current and better efficiency in the red part of the spectrum. Enhanced fluorescence detection efficiency enabled image collection using fast scanning (1 − 5 μ*s*/*pixel*). This enhances further the signal-to-noise-ratio by avoiding fluorescence loss due to triplet state formation, enabling fluorescence imaging with single-molecule sensitivity. In addition, low laser intensities (150- 750 μ*W*) could be applied for imaging, significantly reducing the photo-toxicity (Vukojevic et al., 2008).

FCS measurements are performed by recording fluorescence intensity fluctuations in a very small, approximately ellipsoidal observation volume element (OVE) (about 0.2*μm* wide and 1μ*m* long) that is generated in trachea epithelial cells by focusing the laser light through the microscope objective and by collecting the fluorescence light through the same objective using a pinhole in front of the detector to block out-of-focus light. The fluorescence intensity fluctuations, caused by fluorescently labeled molecules passing through the OVE are analyzed using temporal autocorrelation analysis.

In temporal autocorrelation analysis we first derive the autocorrelation function *G*(*τ*):

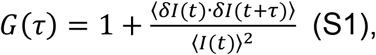

where *δI*(*t*) = *I*(*t*) − ⟨*I*(*t*)⟩ is the deviation from the mean intensity at time *t* and *δI*(*t* + *τ*) = *I*(*t* + *τ*) – ⟨*I* (*t*)⟩ is the deviation from the mean intensity at time *t* + *τ*. For further analysis, an autocorrelation curve is derived by plotting *G*(*τ*) as a function of the lag time, i.e. the autocorrelation time *τ*.

To derive information about molecular numbers and their corresponding diffusion time, the experimentally obtained autocorrelation curves are compared to autocorrelation functions derived for different model systems. A model describing free three dimensional (3D) diffusion of two components and triplet formation was used in this study:

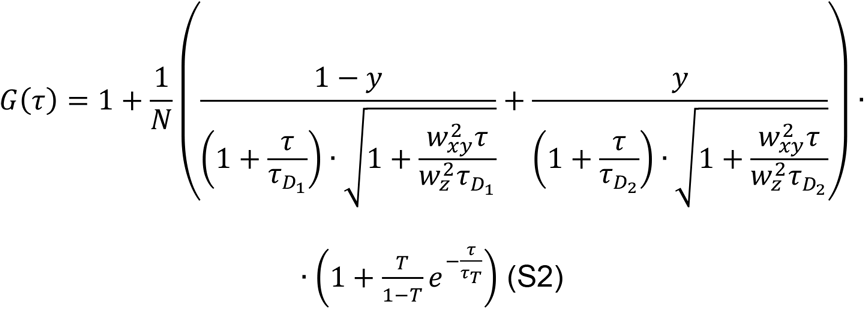

In the above equation, *N* is the average number of molecules in the OVE; *y* is the fraction of the slowly moving Yki-GFP molecules; 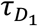 is the diffusion time of the free Yki-GFP molecules; 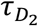 is the diffusion time of Yki-GFP molecules undergoing interactions with the DNA; *W*_*xy*_ and *W*_*z*_ are radial and axial parameters, respectively, related to spatial properties of the OVE; T is the average equilibrium fraction of molecules in the triplet state; and *τ*_*T*_ the triplet correlation time related to rate constants for intersystem crossing and the triplet decay. Spatial properties of the detection volume, represented by the square of the ratio of the axial and radial parameters 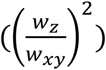, are determined in calibration measurements performed using a solution of Rhodamine 6G for which the diffusion coefficient (D) is known to be *D*_*Rh*6*G*_ = 4.1 ∙ 10^−10^ *m*^2^*s*^−1^ (Muller et al., 2008). The diffusion time, *τ*_*D*_, measured by FCS, is related to the translation diffusion coefficient *D* by:

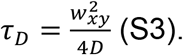

To establish that Yki molecules diffusing through the OVE are the underlying cause of the recorded fluorescence intensity fluctuations, we plotted the characteristic decay times *τ*_*D*1_ and *τ*_*D*2_, obtained by FCS, as a function of the total concentration of Yki molecules. We observed that both characteristic decay times remain stable for increasing total concentration of Yki molecules, signifying that the underlying process triggering the fluorescence intensity fluctuations is diffusion of fluorescent Yki molecules through the OVE (which should be independent of the total concentration of Yki molecules).

### Supplement 3. Sequences for the generation of the CRISPR knock-in line expressing an N-terminal tagged form of Yki (called mKate2-Yki)

The following sequences were used:

a) yki gRNA: 5’-ccttcttcacgcccccggcgccc-3’

b) yki gRNA modified sequence in the donor vector: 5’-cgttttttaccccgcccgccccg-3’

c) donor vector for Yki Knockin with dsRed excisable cassette:

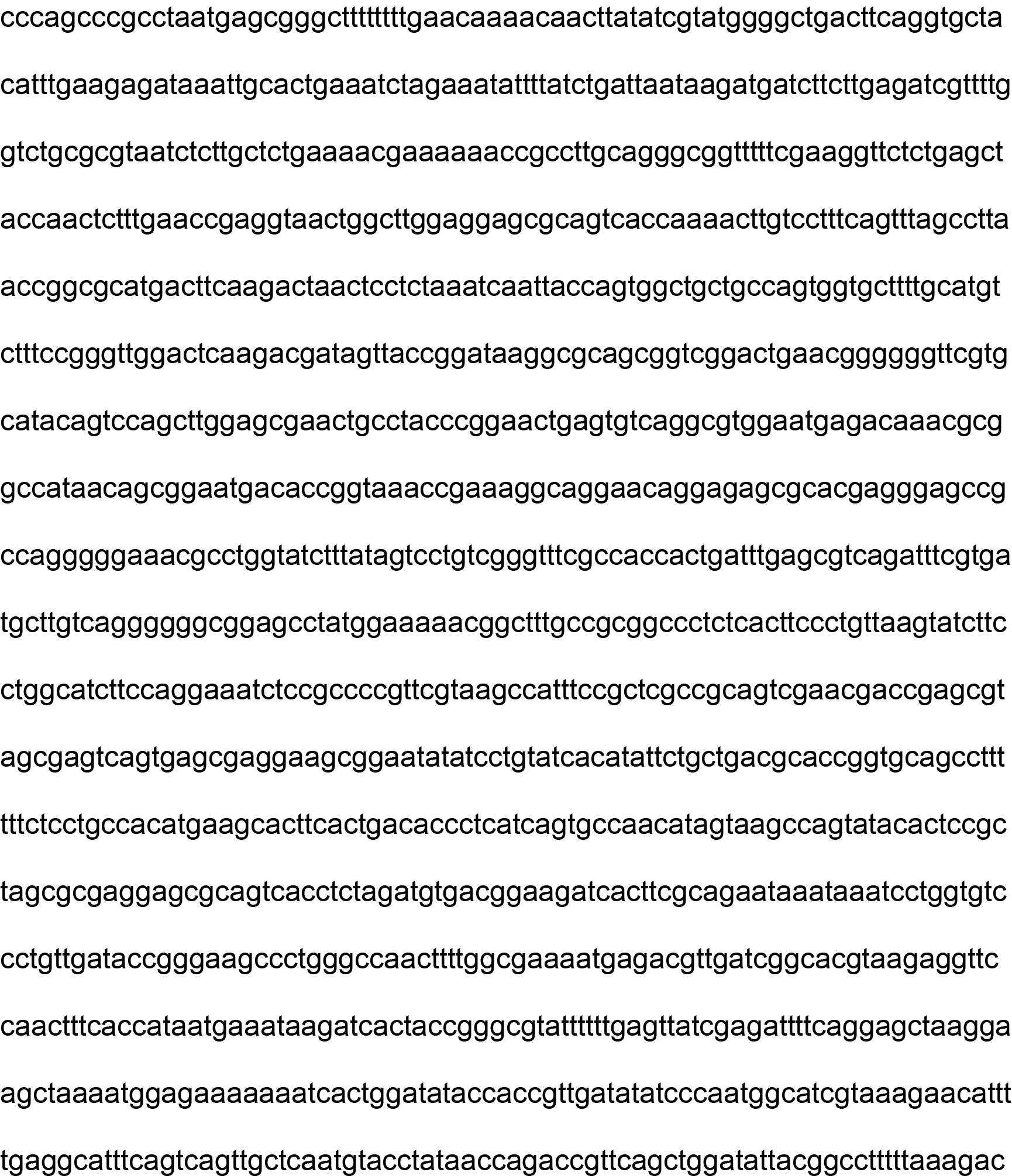

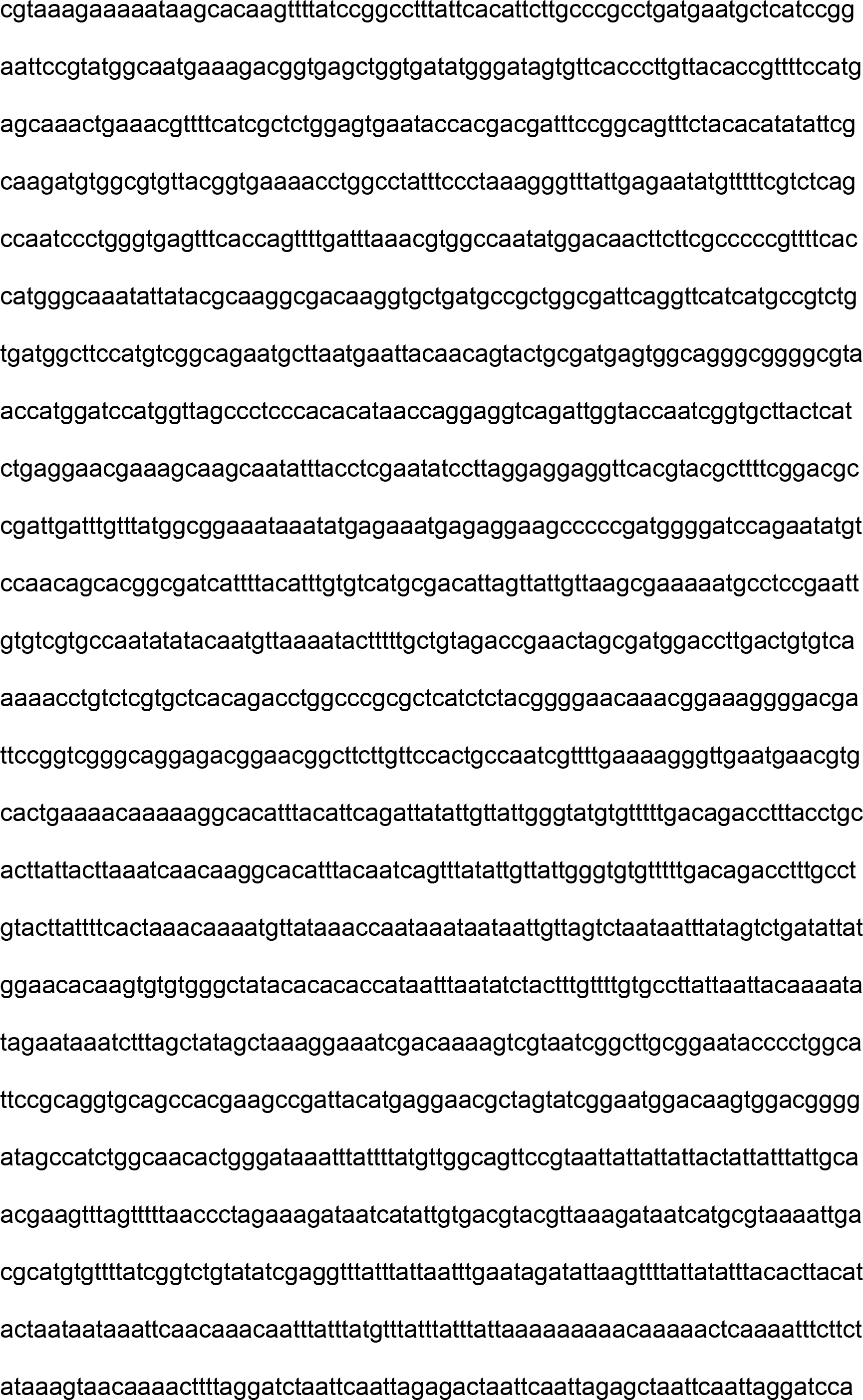

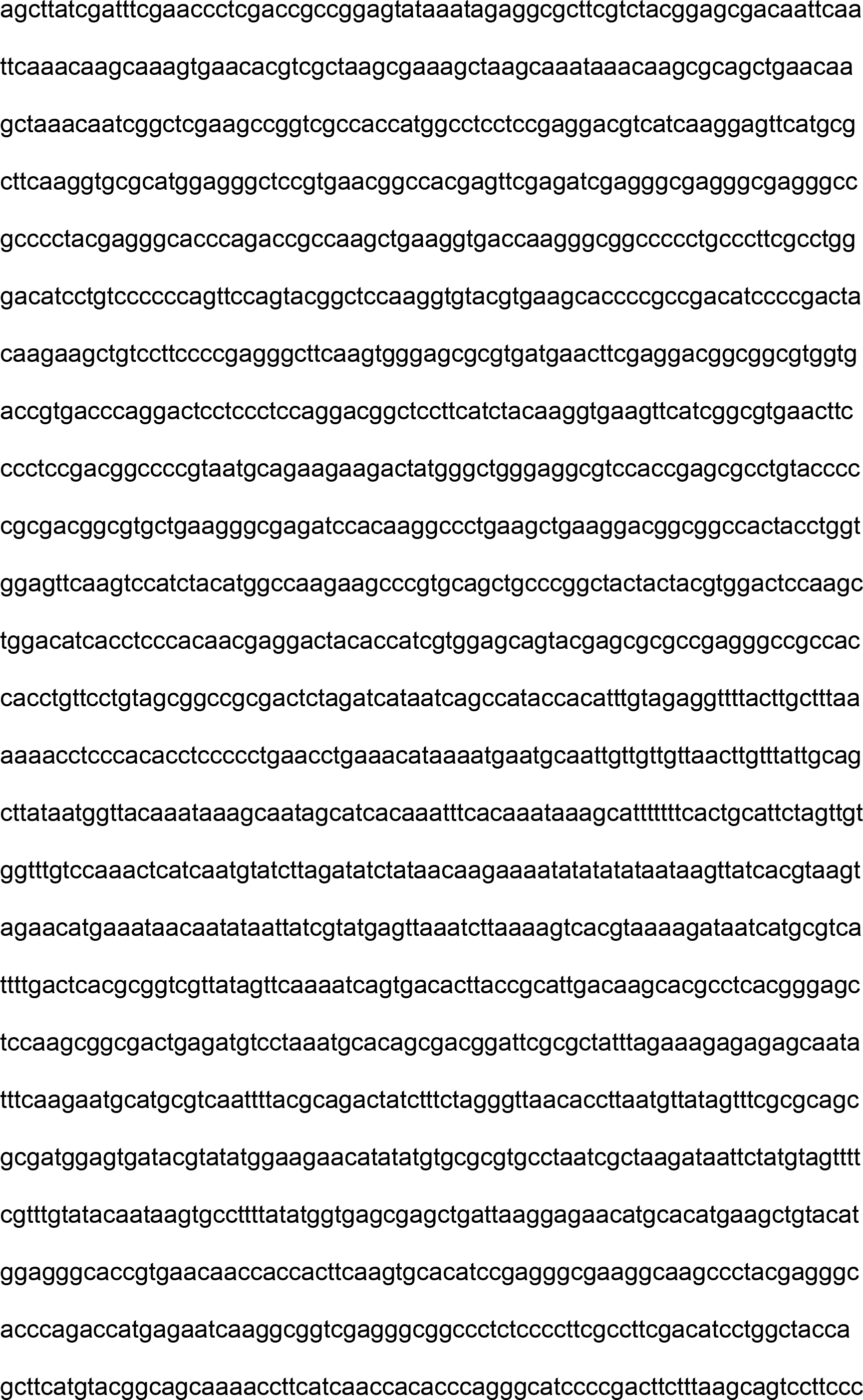

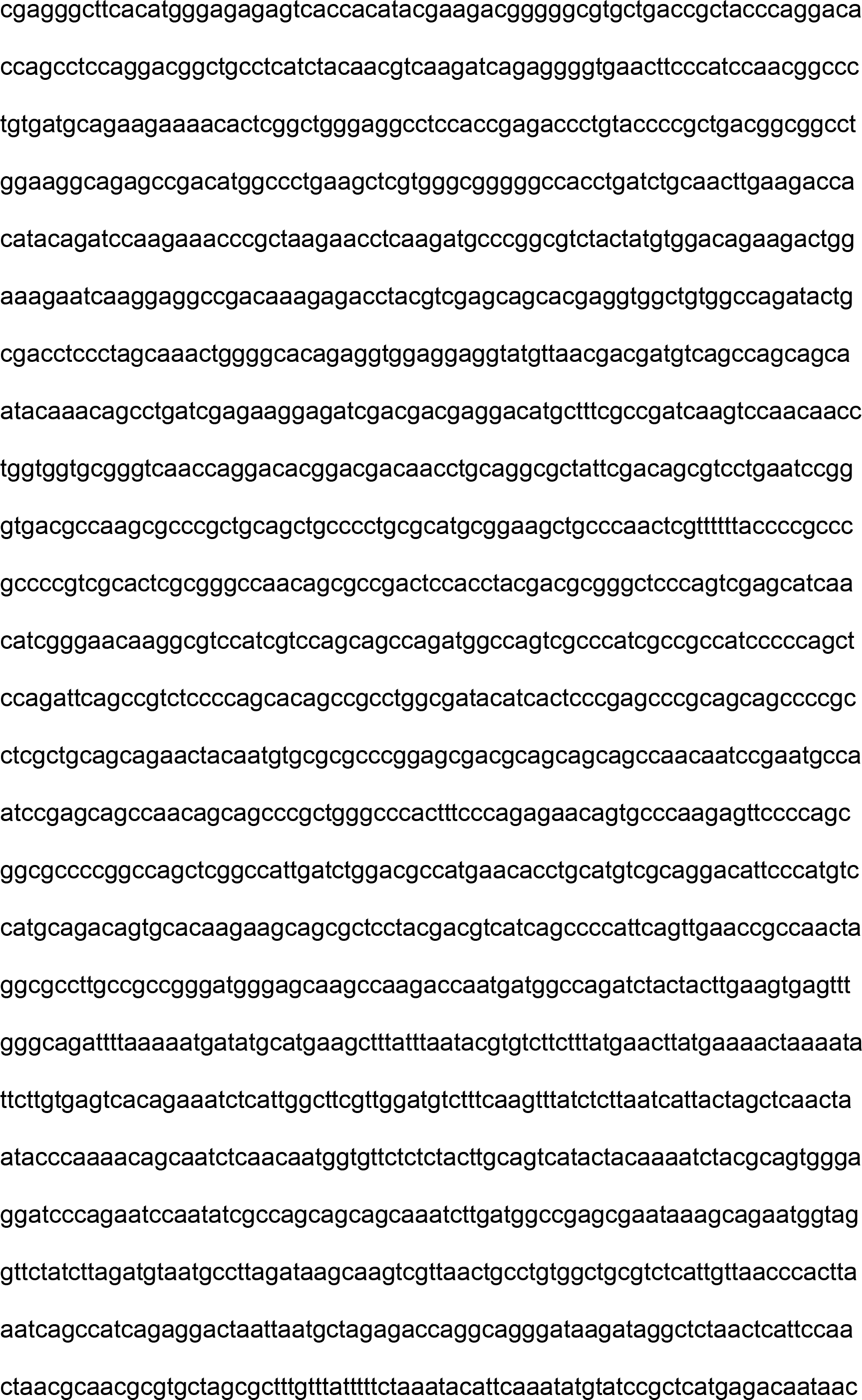

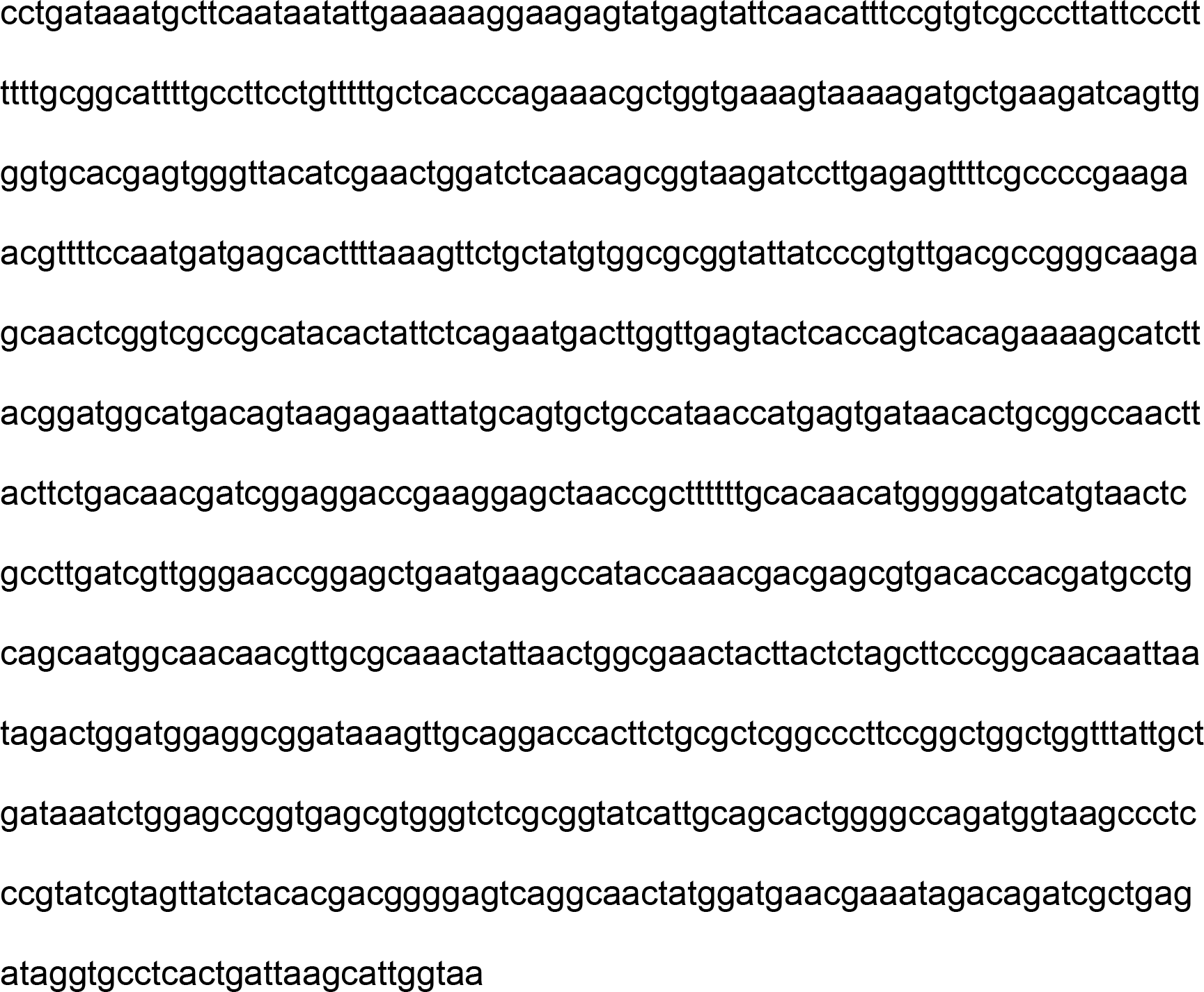

Author contributions: DKP, VT and KS, designed and performed the experiments, analyzed the data, prepared figures; PT, CS and EK supervised the project; KS conceived the project and wrote the manuscript with critical input from all authors; all authors critically reviewed the manuscript and approved it for submission.

